# Successful forecasting of harmful cyanobacteria blooms with high frequency lake data

**DOI:** 10.1101/674325

**Authors:** M.J. Kehoe, B.P. Ingalls, J.J. Venkiteswaran, H.M. Baulch

**Affiliations:** School of Environment and Sustainability and Global Institute for Water Security. University of Saskatchewan. 11 Innovation Blvd. Saskatoon SK, Canada S7N 3H5.; Department of Applied Mathematics. University of Waterloo. 200 University Ave. W Waterloo ON, Canada N2L 3G1.; Department of Geography and Environmental Studies. Wilfrid Laurier University. 75 University Avenue West, Waterloo ON, Canada N2L 3C5.; School of Environment and Sustainability and Global Institute for Water Security. 11 Innovation Blvd. Saskatoon SK, Canada S7N 3H5.

**Keywords:** cyanobacterial bloom, forecasting, high frequency, sensor

## Abstract

Cyanobacterial blooms are causing increasing issues across the globe. Bloom forecasting can facilitate adaptation to blooms. Most bloom forecasting models depend on weekly or fortnightly sampling, but these sparse measurements can miss important dynamics. Here we develop forecasting models from five years of high frequency summer monitoring in a shallow lake (which serves as an important regional water supply). A suite of models were calibrated to predict cyanobacterial fluorescence (a biomass proxy) using measurements of: cyanobacterial fluorescence, water temperature, light, and wind speed. High temporal autocorrelation contributed to relatively strong predictive power over 1, 4 and 7 day intervals. Higher order derivatives of water temperature helped improve forecasting accuracy. While traditional monitoring and modelling have supported forecasting on longer timescales, we show high frequency monitoring combined with telemetry allows forecasting over timescales of 1 day to 1 week, supporting early warning, enhanced monitoring, and adaptation of water treatment processes.

## Introduction

Blooms of cyanobacteria are an increasing problem around the world (Hoagland et al., 2002; Kotak and Zurawell, 2007) including regions with cooler climates (Pick, 2016, Mantzouki et al., 2018). Nuisance biomass of algae and cyanobacteria can lead to degradation of ecosystem services, loss of property values, and high costs for drinking water treatment (Dodds et al., 2009; Hudnell, 2010). Blooms of cyanobacteria can also lead to issues of unpleasant taste and odour and can have direct impacts on the safety of drinking water supplies by producing a variety of toxins (WHO and Unicef, 2011) which also pose health risks for swimmers and boaters (Dodds et al., 2009; Hudnell, 2010). In light of growing bloom risk and slow progress towards bloom mitigation, there is a need to develop adaptation strategies. These adaptation strategies may include improved risk communications, increased monitoring, altered drinking water treatment, and bloom forecasting.

Several groups can make use of bloom forecasts: water treatment plant operators, recreational users, and regulatory agencies responsible for toxin testing. Water treatment plants benefit from bloom forecasts providing advance warning of necessary changes in treatment processes to adapt to bloom conditions, such as addition of activated carbon or modification of coagulant dose. Health agencies and water treatment plant operators can identify when they will need to test for toxins, and recreational users can avoid bloom-affected lakes. Forecasting is also of major benefit to scientists, who may ‘chase’ blooms - but rarely capture antecedent conditions.

While bloom forecasting is well-advanced in the oceans (Seltenrich, 2014), it remains an underdeveloped area in freshwaters. Bloom forecasts have been operationalized only in a small number of very well-studied ecosystems, such as Lake Erie (Obenour et al., 2014) and Lake Taihu (Kong et al., 2009). Forecasting approaches differ in the specific variables being forecast, the methods employed, and the forecasting period. Forecasts of variables such as cyanobacterial cell counts or cyanobacterial biovolume have been developed using regression-based approaches – predicting two weeks into the future (Onderka, 2007; Persaud et al., 2015). Cyanotoxin-forecasting approaches have included genetic algorithms and artificial neural networks (Chan et al., 2007; Fernández et al., 2013). The density of specific cyanobacterial taxa can also be predicted using approaches such as hybrid evolutionary algorithms (Welk et al., 2008) on relatively short timescales (5-7 days) using decadal or longer datasets from discrete sampling events (Welk et al., 2008). Hydrodynamic models and coupled hydrodynamic-algal biomass models have been useful in understanding bloom development and transport (Wynne et al., 2013; Li et al., 2014).

The growing availability of high frequency monitoring data provides new opportunities for bloom forecasting (Hamilton et al., 2015). Compared with traditional weekly or fortnightly sampling, high frequency monitoring is much better matched to understanding and predicting blooms, due to the short temporal scales over which blooms develop and disappear. Here we address the question: How well can we predict cyanobacteria blooms using high frequency monitoring data? We assess a range of approaches to bloom forecasting and develop recommendations for simple forecasting tools based on insights from a shallow polymictic lake which represents a crucial regional drinking water supply in the Canadian prairies.

## Methods

### Study site

Buffalo Pound Lake is a long, shallow lake (35 km long, 1 km wide, mean depth approximately 3 m, maximum depth 5.6 m) which experiences frequent mixing due to strong winds and nighttime convection. Transient stratification events occur in summer when winds are low or blowing across the narrow axis of the lake. Buffalo Pound is eutrophic (Hammer, 1978), and is subject to frequent blooms affecting operations of the water treatment plant that supplies drinking water to the cities of Regina and Moose Jaw, Saskatchewan (Kehoe et al., 2015).

### Dataset

The dataset consists of five years of high frequency monitoring data from Buffalo Pound Lake during the open water season. Data was collected from a single buoy positioned in the middle of the lake near the water treatment plant intake at 50.58611 N 105.38472 W. The dataset describes phycocyanin relative fluorescence measured at 0.82 m (YSI 6600 Sonde), temperature at 0.45 m, 0.77 m, 0.82 m, 1.23 m, 2.18 m, 2.85 m, 3.18 m (combination of YSI 6600, YSI 6920 and Nexsens T-Node FR sensors), wind speed measured with a Vaisala WXT536 sensor (1.66 m above the water surface), and photosynthetically active radiation (PAR) measured at the top of the buoy (1.06 m above the water surface) with a LiCor L-I190 2 pi sensor.

The following variables were used in forecast models: phycocyanin relative fluorescence, water column stability, mean water column temperature, wind speed and surface PAR. Phycocyanin fluorescence is generated by the photosynthetic pigment phycocyanin which is specific to cyanobacteria. Details on the relationship between observed phycocyanin fluorescence and observed cyanobacterial biomass can be found in the Appendix. Water column stability was calculated using the Schmidt stability index function ‘schmidt.stability’ provided with the R package ‘rLakeAnalyzer’ (Winslow et al., 2018). The ‘schmidt.stability’ function calculates water column stability using all temperature measurements (i.e. all depths) as well as bathymetry data of cross sectional areas and corresponding depths which was derived from digital elevation data using ArcGIS. Raw fluorescence data were preprocessed to reduce noise using the ‘signal’ (signal developers, 2014) R package. First, all variables were low-pass filtered with a Butterworth filter – order: 5 frequency: day^-1^. Next a 3^rd^ order Savitsky-Golay filter of order p=3 and length n=145 was applied to extract higher derivatives – acceleration and jerk – of water temperature.

### Modelling

The following predictors were used to forecast future phycocyanin fluorescence at time horizons of 1, 4 and 7 days: current phycocyanin fluorescence, wind speed, water temperature, incident photosynthetically active radiation (PAR), water column stability, 1st derivative of water temperature (*TD*), 2nd derivative of water temperature (*TA*), 3rd derivative of water temperature (*TJ*).

Our model design begins with the exponential growth equation:

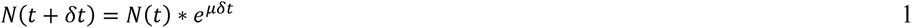

Where *N* is the abundance of phycocyanin fluorescence, *t* is time, *δt* is the size of the time step, and *µ* is specific growth rate.

Taking the natural log of the above yields a form amenable to our purposes:

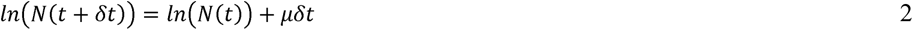

Five statistical and machine learning models were used to model the function above (Eqn 2.): boosted decision tree (bdt), decision tree (dt), linear regression (lm), boosted linear regression (blm) and a non-linear model (nlm) we outline below. A boosted decision tree is a higher order model made up of decision trees and a boosted linear regression is an ensemble of linear regressions. Both boosted decision trees and boosted linear regressions are capable of representing more complex non-linear relationship than the linear regression based models. By including both the simple forms and the more complex forms we can determine how much model complexity is warranted for this forecasting problem.

A non-linear model (nlm) of the following form was also fitted, following the same procedure as the statistical and machine learning models:

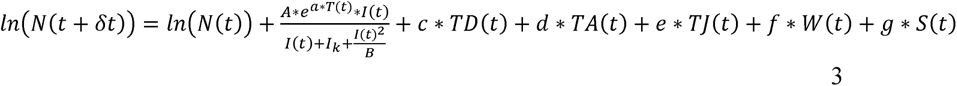

where *TD*(*t*) denotes the 1^st^ derivative of water temperature, *TA*(*t*) denotes the 2^nd^ derivative of water temperature, *TJ*(*t*) denotes the 3^rd^ derivative of water temperature, *w*(*t*) denotes wind, and *S*(*t*) denotes water column stability. The term 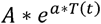 captures the exponential change in maximum growth rate with temperature. PAR is denoted by *I*(*t*). The term 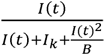 allows the possibility -depending on estimate parameters- of light limitation and inhibition due to PAR. The symbols *a, A, B, c, d, e, f, g*, are parameters to be estimated.

All analysis was carried out with R (R Core Team, 2017). For linear modelling and non-linear modelling the functions ‘lm’ and ‘nls’ functions from the base package were used (R Core Team, 2017). Boosted linear regression and boosted decision tree were constructed with the “xgboost” package (Chen et al., 2018), and decision trees with ‘rpart’(Therneau et al., 2017). Plot construction was carried out with the base plotting options and ‘ggplot’ (Wickham, 2016).

The models were assessed via cross-validation. Models were calibrated on five different subsets of the total of five years, each of which omitted a different year for model validation. Models were calibrated for each of 1, 4 and 7 day forecast horizons. Model performance was assessed using three different metrics applied to the out-of-calibration year of data. First, Pearson correlation (ρ), commonly called ‘forecast skill’ was used to test the ability to predict dynamics. Second, the area under the curve (AUC) for the receiver operator curve was used to assess the model ability to predict extreme events - here defined as the 0.75 quantile or above of phycocyanin fluorescence. Finally, range normalized root mean square error (nRMSE) was used to test model fit – i.e., the deviation between predicted and observed values:

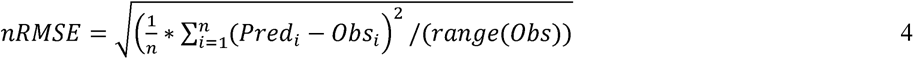

Where *Pred* are predicted values and *Obs* are observed values and *n* is number of observations in a given year. *range*(*obs*) are the range of observations in a year. Normalization by this term allows comparison between years.

Model performance metrics were all compared to that of a naïve model that simply predicts no change – the future measurement will be identical to the most recent measurement.

## Results & Discussion

Reasonably accurate seasonal cyanobacterial bloom forecasts are now achievable, and have been operationalized in several bloom affected ecosystems (Kong et al., 2009; Obenour et al., 2014). Here, we demonstrate the accuracy of short-term cyanobacterial bloom forecasts informed by autonomous sensors - which show relatively strong capacity to predict cyanobacterial blooms on short (1-7 day) timescales with limited input data (Tables 1-3). Sensor data showed a reasonable relationship with observed cyanobacterial biomass, despite known limitations of the technology (Hodges et al., 2018), and small differences in depth and location of the water treatment plant intake and sensor mooring (Appendix Figure 1). We note that 14 day forecasts were uniformly poor, particularly for 2014 and 2015 (Tables 1-3). In all models, the high degree of autocorrelation in the phycocyanin fluorescence data lead to current phycocyanin as the strongest model predictor, with water temperature and changes in temperature higher derivatives, PAR and wind providing added insights. This high degree of autocorrelation is unsurprising because changes in phytoplankton biomass are biomass dependent - yet suggest that very simple approaches to forecasting may be effective in some lakes.

**Table 1.** Validation performance of all models predicting phycocyanin fluorescence 1, 4, 7 and 14 days ahead as measured by the range normalized root mean square error (nRMSE). Light greyed in cells denote model/s with nRMSE less than the nRMSE of the naïve forecast model. Darker grey squares denote the model/s with the lowest nRMSE compared to the naïve forecast model. The individual models tested were decision tree (dt), boosted decision tree (bdt), linear model (lm), boosted linear model (blm), nonlinear model (nlm). Note that nRMSE has no units as it is root mean square error normalized to the range of observations.

| Validation data set | nRMSE-dt | nRMSE-bdt | nRMSE-lm | nRMSE-blm | nRMSE-nlm | nRMSE naïve model<br>$N(t)=N(t-1)$ |
| --- | --- | --- | --- | --- | --- | --- |
| **2014** | 0.11 | 0.12 | 0.09 | 0.09 | 0.09 | 0.09 |
| **2015** | 0.10 | 0.10 | 0.08 | 0.08 | 0.08 | 0.09 |
| **2016** | 0.07 | 0.09 | 0.05 | 0.05 | 0.05 | 0.05 |
| **2017** | 0.06 | 0.07 | 0.05 | 0.05 | 0.04 | 0.04 |
| **2018** | 0.07 | 0.08 | 0.06 | 0.06 | 0.06 | 0.06 |

Four day forecast
| Validation data set | nRMSE-dt | nRMSE-bdt | nRMSE-lm | nRMSE-blm | nRMSE-nlm | nRMSE naïve model<br>$N(t)=N(t-1)$ |
| --- | --- | --- | --- | --- | --- | --- |
| **2014** | 0.26 | 0.28 | 0.24 | 0.24 | 0.26 | 0.28 |
| **2015** | 0.23 | 0.21 | 0.22 | 0.22 | 0.22 | 0.24 |
| **2016** | 0.11 | 0.15 | 0.09 | 0.09 | 0.08 | 0.09 |
| **2017** | 0.13 | 0.13 | 0.11 | 0.11 | 0.09 | 0.10 |
| **2018** | 0.12 | 0.13 | 0.10 | 0.10 | 0.10 | 0.11 |

Seven day forecast
| Validation data set | nRMSE-dt | nRMSE-bdt | nRMSE-lm | nRMSE-blm | nRMSE-nlm | nRMSE naïve model $N(t)=N(t-1)$ |
| --- | --- | --- | --- | --- | --- | --- |
| **2014** | 0.46 | 0.47 | 0.47 | 0.47 | 0.54 | 0.58 |
| **2015** | 0.26 | 0.22 | 0.24 | 0.24 | 0.24 | 0.29 |
| **2016** | 0.18 | 0.20 | 0.13 | 0.12 | 0.12 | 0.13 |
| **2017** | 0.16 | 0.14 | 0.13 | 0.14 | 0.11 | 0.13 |
| **2018** | 0.15 | 0.15 | 0.13 | 0.13 | 0.14 | 0.17 |
| Fourteen day forecast |  |  |  |  |  |  |
| Validation data set | nRMSE-dt | nRMSE-bdt | nRMSE-lm | nRMSE-blm | nRMSE-nlm | nRMSE naïve model $N(t)=N(t-1)$ |
| **2014** | 0.63 | 0.56 | 0.65 | 0.63 | 0.80 | 0.84 |
| **2015** | 0.25 | 0.23 | 0.23 | 0.22 | 0.24 | 0.27 |
| **2016** | 0.25 | 0.27 | 0.18 | 0.18 | 0.18 | 0.20 |
| **2017** | 0.19 | 0.18 | 0.18 | 0.19 | 0.14 | 0.19 |
| **2018** | 0.23 | 0.24 | 0.20 | 0.20 | 0.23 | 0.31 |

**Table 2.** Validation performance of all models predicting phycocyanin fluorescence 1, 4 and 7 days ahead as measured by the Pearson correlation coefficient (ρ). Light greyed in cells denote model/s with Pearson correlation greater than the Ρearson correlation of the naïve forecast model. Darker grey squares denote the models with the highest Pearson correlation compared to the naïve forecast model.

| Validation data set | $\rho$ -dt | $\rho$ -bdt | $\rho$ -lm | $\rho$ -blm | $\rho$ - nlm | $\rho$ naïve model<br>$N(t)=N(t-1)$ |
| --- | --- | --- | --- | --- | --- | --- |
| **2014** | 0.87 | 0.89 | 0.91 | 0.91 | 0.91 | 0.92 |
| **2015** | 0.87 | 0.87 | 0.90 | 0.90 | 0.90 | 0.89 |
| **2016** | 0.97 | 0.98 | 0.99 | 0.99 | 0.99 | 0.98 |
| **2017** | 0.98 | 0.98 | 0.99 | 0.99 | 0.99 | 0.99 |
| **2018** | 0.94 | 0.95 | 0.96 | 0.96 | 0.96 | 0.95 |

Four day forecast
| Validation data set | $\rho$ -dt | $\rho$ -bdt | $\rho$ -lm | $\rho$ -blm | $\rho$ -nlm | $\rho$ naïve model<br>$N(t)=N(t-1)$ |
| --- | --- | --- | --- | --- | --- | --- |
| **2014** | 0.45 | 0.45 | 0.53 | 0.53 | 0.55 | 0.55 |
| **2015** | 0.43 | 0.45 | 0.44 | 0.44 | 0.44 | 0.44 |
| **2016** | 0.91 | 0.91 | 0.95 | 0.96 | 0.96 | 0.96 |
| **2017** | 0.91 | 0.91 | 0.94 | 0.94 | 0.94 | 0.93 |
| **2018** | 0.80 | 0.85 | 0.85 | 0.85 | 0.85 | 0.85 |

Seven day forecast
| Validation data set | $\rho$ -dt | $\rho$ -bdt | $\rho$ -lm | $\rho$ -blm | $\rho$ - nlm | $\rho$ naïve model $N(t)=N(t-1)$ |
| --- | --- | --- | --- | --- | --- | --- |
| **2014** | -0.21 | -0.15 | -0.26 | -0.26 | -0.24 | -0.24 |
| **2015** | 0.34 | 0.40 | 0.42 | 0.41 | 0.41 | 0.38 |
| **2016** | 0.84 | 0.85 | 0.93 | 0.94 | 0.93 | 0.94 |
| **2017** | 0.88 | 0.89 | 0.91 | 0.90 | 0.92 | 0.89 |
| **2018** | 0.68 | 0.68 | 0.74 | 0.73 | 0.73 | 0.71 |

| Validation<br>data set | $\rho$ -dt | $\rho$ -bdt | $\rho$ -lm | $\rho$ -blm | $\rho$ - nlm | $\rho$ naïve model $N(t)=N(t-1)$ |
| --- | --- | --- | --- | --- | --- | --- |
| **2014** | -0.56 | -0.66 | -0.69 | -0.69 | -0.66 | -0.66 |
| **2015** | 0.33 | 0.32 | 0.50 | 0.49 | 0.47 | 0.46 |
| **2016** | 0.81 | 0.76 | 0.90 | 0.91 | 0.90 | 0.92 |
| **2017** | 0.81 | 0.79 | 0.83 | 0.81 | 0.85 | 0.78 |
| **2018** | 0.44 | 0.44 | 0.47 | 0.44 | 0.44 | 0.37 |

**Table 3.** Validation performance of all models predicting 1, 4 and 7 days ahead as measured by the area under the curve (AUC) for the the receiver operating characteristics curve using the 0.75 quantile of phycocyanin fluorescence as the cutoff for conditions deemed ‘bloom’. Light greyed in cells denote model/s with AUC greater than the AUC of the naïve forecast model. Darker grey squares denote the model/s with the highest AUC compared to the naïve forecast model. The individual models tested were decision tree (dt), boosted decision tree (bdt), linear model (lm), boosted linear model (blm), nonlinear model (nlm).

| Validation data set | AUC -dt | AUC -bdt | AUC -lm | AUC -blm | AUC-nlm | AUC naïve model<br>N(t)=N(t-1) |
| --- | --- | --- | --- | --- | --- | --- |
| **2014** | 0.90 | 0.88 | 0.92 | 0.92 | 0.92 | 0.92 |
| **2015** | 0.92 | 0.94 | 0.95 | 0.95 | 0.95 | 0.95 |
| **2016** | 0.85 | 0.95 | 0.96 | 0.96 | 0.96 | 0.96 |
| **2017** | 0.99 | 0.99 | 0.99 | 0.99 | 0.99 | 0.99 |
| **2018** | 0.92 | 0.98 | 0.98 | 0.98 | 0.98 | 0.98 |

Four day forecast
| Validation data set | AUC -dt | AUC -bdt | AUC -lm | AUC -blm | AUC-nlm | AUC naïve model<br>N(t)=N(t-1) |
| --- | --- | --- | --- | --- | --- | --- |
| **2014** | 0.52 | 0.57 | 0.56 | 0.63 | 0.56 | 0.56 |
| **2015** | 0.70 | 0.73 | 0.74 | 0.74 | 0.73 | 0.73 |
| **2016** | 0.78 | 0.90 | 0.88 | 0.86 | 0.85 | 0.83 |
| **2017** | 0.93 | 0.97 | 0.97 | 0.96 | 0.97 | 0.96 |
| **2018** | 0.82 | 0.85 | 0.88 | 0.87 | 0.87 | 0.86 |

Seven day forecast
| Validation data set | AUC -dt | AUC -bdt | AUC -lm | AUC -blm | AUC-nlm | AUC naïve model<br>$N(t)=N(t-1)$ |
| --- | --- | --- | --- | --- | --- | --- |
| **2014** | 0.78 | 0.81 | 0.81 | 0.81 | 0.81 | 0.80 |
| **2015** | 0.67 | 0.73 | 0.74 | 0.73 | 0.74 | 0.71 |
| **2016** | 0.84 | 0.89 | 0.87 | 0.85 | 0.87 | 0.81 |
| **2017** | 0.91 | 0.94 | 0.95 | 0.95 | 0.96 | 0.94 |
| **2018** | 0.87 | 0.85 | 0.79 | 0.79 | 0.79 | 0.78 |

Fourteen day forecast
| Validation data set | AUC -dt | AUC -bdt | AUC -lm | AUC -blm | AUC-nlm | AUC naïve model<br>$N(t)=N(t-1)$ |
| --- | --- | --- | --- | --- | --- | --- |
| **2014** | 0.73 | 0.82 | 0.84 | 0.84 | 0.82 | 0.84 |
| **2015** | 0.65 | 0.70 | 0.77 | 0.75 | 0.75 | 0.73 |
| **2016** | 0.78 | 0.78 | 0.84 | 0.80 | 0.85 | 0.72 |
| **2017** | 0.88 | 0.89 | 0.87 | 0.85 | 0.87 | 0.81 |
| **2018** | 0.59 | 0.62 | 0.61 | 0.58 | 0.58 | 0.54 |

The performance of the linear and non-linear regression models improved in comparison to the naïve model (which assumes a correlation of r=1) as the forecasting time scale increased. The naïve model’s performance deteriorates over longer horizons because autocorrelation is reduced at longer time scales. We anticipate that accurate longer term forecasts (>7 days) may not be achievable without including a description of the key mechanisms affecting bloom risk. Here, we see that 14 day forecasts were poor for all years and approaches (Tables 1-3 and see Appendix plots A16-A20), although they did often out-perform the naïve model. Mechanistic approaches to bloom risk involve added challenges, for example, associated with forecasting factors such as weather or nutrient runoff. Nonetheless, strong relationships between seasonal or annual nutrient loads and bloom severity (e.g., Binding et al. 2018) suggest that parsimonious approaches to seasonal forecasts in shallow lakes may be feasible.

The five bloom years studied here showed major differences in dynamics and timing (Figure 1); yet cross validation results were promising. However, models were not able to forecast abrupt increases in biomass (e.g. Figure 2) and most of our models failed to forecast a spike in biomass in 2017 which was associated with high light and high temperature (see Appendix plots A5, A10, A14, A19). Late season dynamics are also poorly forecasted in some cases (e.g. Appendix plot A15). We think this is because five years of data is still a relatively small dataset to characterize bloom behavior in Buffalo Pound - given high interannual variation in bloom phenology.

**Figure 1.**
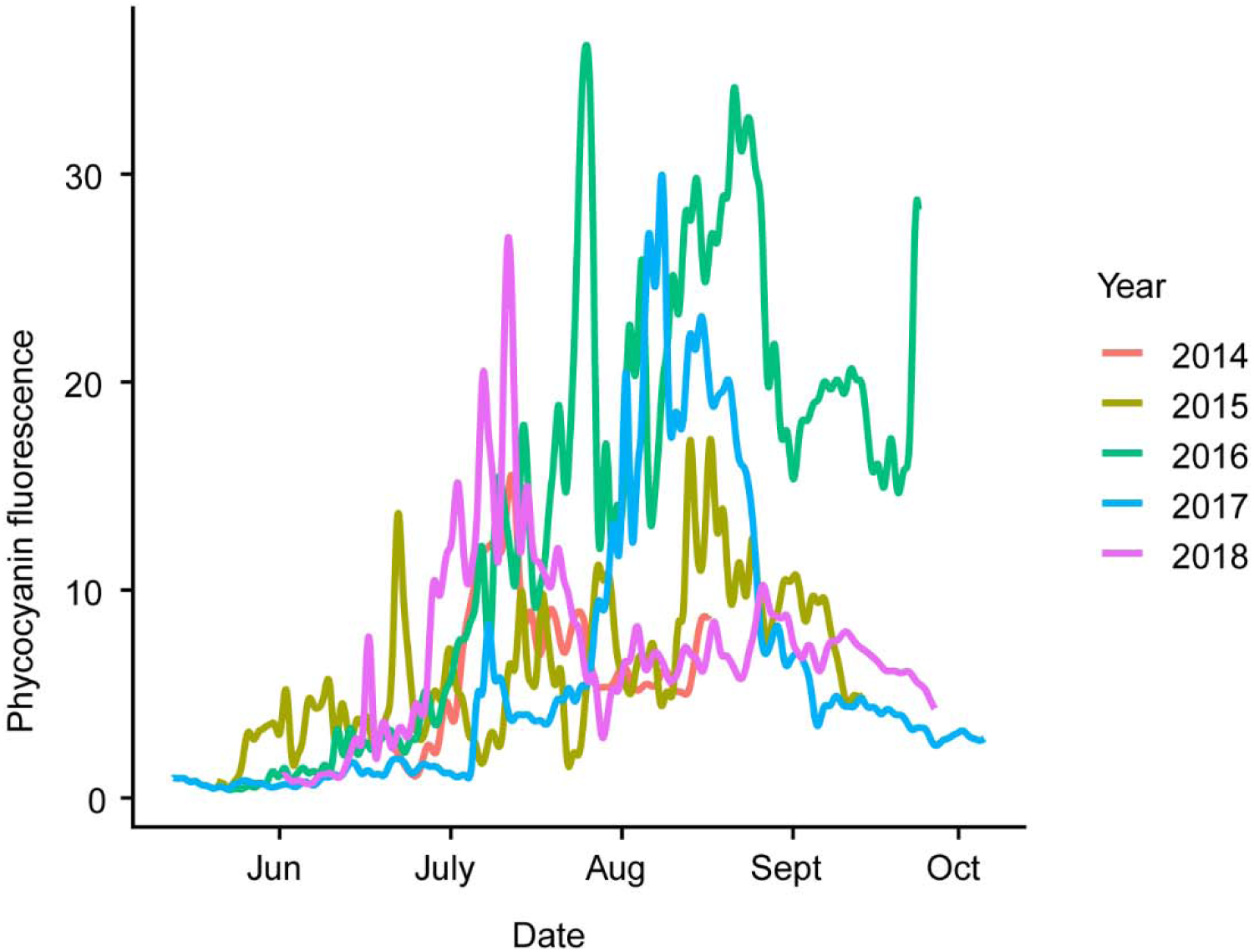
Phycocyanin fluorescence for year of sampling 2014-2018 inclusive. The unit for phycocyanin fluorescence is relative fluorescence units (RFU).

**Figure 2.**
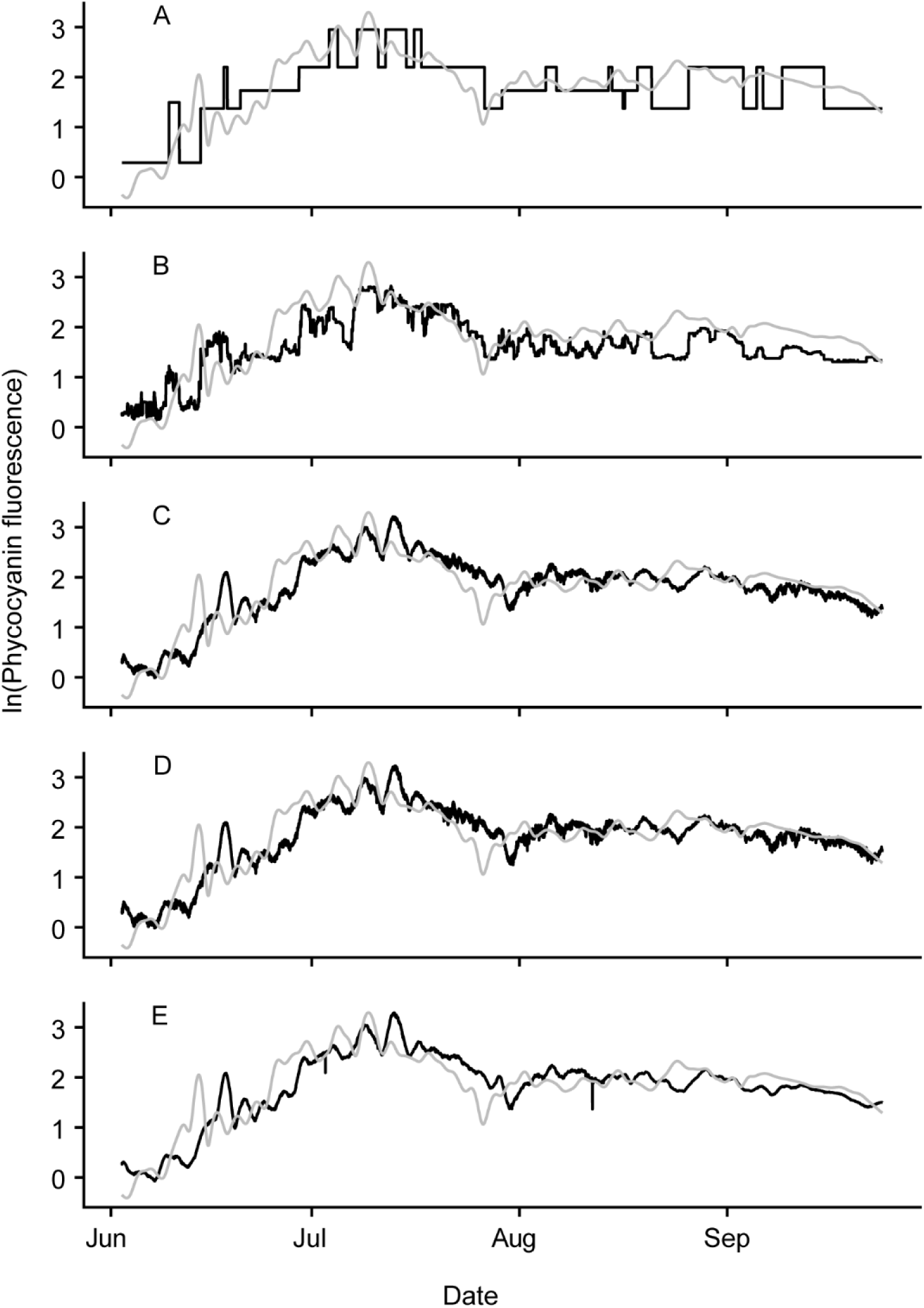
Plots of model performance for 5 different types of models at 4 day time horizon for 2018. A) decision tree (dt), B) boosted decision tree (bdt), C) linear regression (lm), D) boosted linear regression, E) non-linear model. Grey line is observed while black line is forecasted. Plots for other years and forecast horizons can be found in the Appendix. The unit for phycocyanin fluorescence is relative fluorescence unit (RFU).

The linear regression model makes the best choice for operationalization due to its simplicity, reliability, and overall strong performance in the forecast metrics. While the apparent adequacy of linear regression models should offer caution for the use of more complex models like boosted decision trees or boosted regression, we note that the more complex models occasionally showed improvements in the AUC metric at 7 and 14 day forecasts (see Tables 1-3 and Appendix). We also note this work is based only on 5 summers of high frequency data, hence may not provide sufficient data to see the potential strengths of machine learning methods like boosted decision trees, which are notably data intensive and are most effective when data contains significant and persistent non-linear relationships (e.g. Thomas et al., 2018).

Modelling studies like this provide benchmarks against which more complex process-based models can be compared. Our study is one of the few in algal/cyanobacterial forecasting which reports model validation on unseen testing data and which includes a comparison to a naïve model, approaches we recommend should be standard within bloom forecasting. Naïve models provide context to model performance and provide an indication of forecast difficulty - here demonstrating that parsimonious (naïve) models are remarkably effective. Validation allows assessment of how effectively a model structure generalizes, here again showing relatively good performance despite high interannual variation. We also note that the cyanobacteria bloom community requires benchmark datasets against which different models can be compared.

In summary, short term cyanobacterial bloom forecasting on the order of a day to a week seems feasible based on simple approaches using high-frequency monitoring data. Our study shows that cyanobacteria (represented here by phycocyanin relative fluorescence) can be highly predictable over short to medium time frames due to autocorrelation of biomass. All the other input factors - such as water temperature, wind speed and light - only slightly improve forecasting ability.

Future work should test multiple approaches across lakes, to better understand which forecasting methods are the most robust. Future work may also benefit from blending statistical and process-based models, for example, integrating physical processes such as advection to enable prediction of surface scums or extend short-term forecasts to longer timescales.

## Acknowledgements

This research was funded by the Canada First Research Excellence Fund (CFREF), and conducted under the Global Water Futures funded project FORMBLOOM (Forecasting tools and mitigation options for diverse bloom-affected lakes). High frequency monitoring data has been supported by instrumentation funded by the Canada Foundation for Innovation and past funding via an NSERC Strategic Project Grant to HMB. Numerous technical staff have been involved in supporting these measurements, including Jay Bauer, Katy Nugent and others. Support of the Saskatchewan Water Security Agency and Buffalo Pound Water Treatment in logistics, and research is gratefully acknowledged as have valuable discussions and learning opportunities via GLEON. Data and code are available at URL (upon publication).

### Appendix A

#### Validation of phycocyanin assay

To assess the reliability of sensor-based measurements in understanding cyanobacterial blooms and associated risk to the source water supply of the Buffalo Pound Water treatment plant, we compared sensor data to direct measurements of cyanobacterial biomass. 20 samples were collected at discrete time points during the buoy deployment period of 2014, 2015, 2016, and 2017. Samples were obtained from the plant intake, which obtains water from near the lake bottom (∼3m depth) a short distance from the buoy site (50° 35’ 2” N, 105° 23’ 13” W). Samples were preserved by adding 1 mL Lugol’s solution to raw water, and concentrated by settling for 24 h in a 100 mL graduated cylinder. Additional Lugol’s solution was added as required to ensure adequate preservation of high biomass samples. Phytoplankton identification and counts were performed following Findlay and Kling 2001 by Plankton R Us (Winnipeg, Manitoba) using the modified Ütermohl technique (Nauwerck 1963) on an inverted microscope using phase contrast illumination at magnifications of 125, 400 and 1200x. We report the cyanobacterial biomass, with wet-weight biomass estimated by approximating the cell volume from dimensions and geometry (Vollenweider, 1968; Rott, 1981), and assuming a specific gravity of 1. The sensor-based fluorescence measurements were correlated to microscopic biomass measurements, with a R^2^ of 0.64 (Figure A1).

#### Model forecasts

Plots were made comparing observed values to forecasted values for each of the 5 models, for each of the 4 time horizons (1, 4, 7, and 14 days), for each of the 5 years. This resulted in 19 plots (Figs. A2 –A20). That is, one for each year minus the plot for 4 day forecasts for 2018 which are presented in the main text. Plots show the reduction in model performance over longer forecasting timespans, with specific model performance metrics reported in Tables 1-3 in the main text.

**Figure A1.**
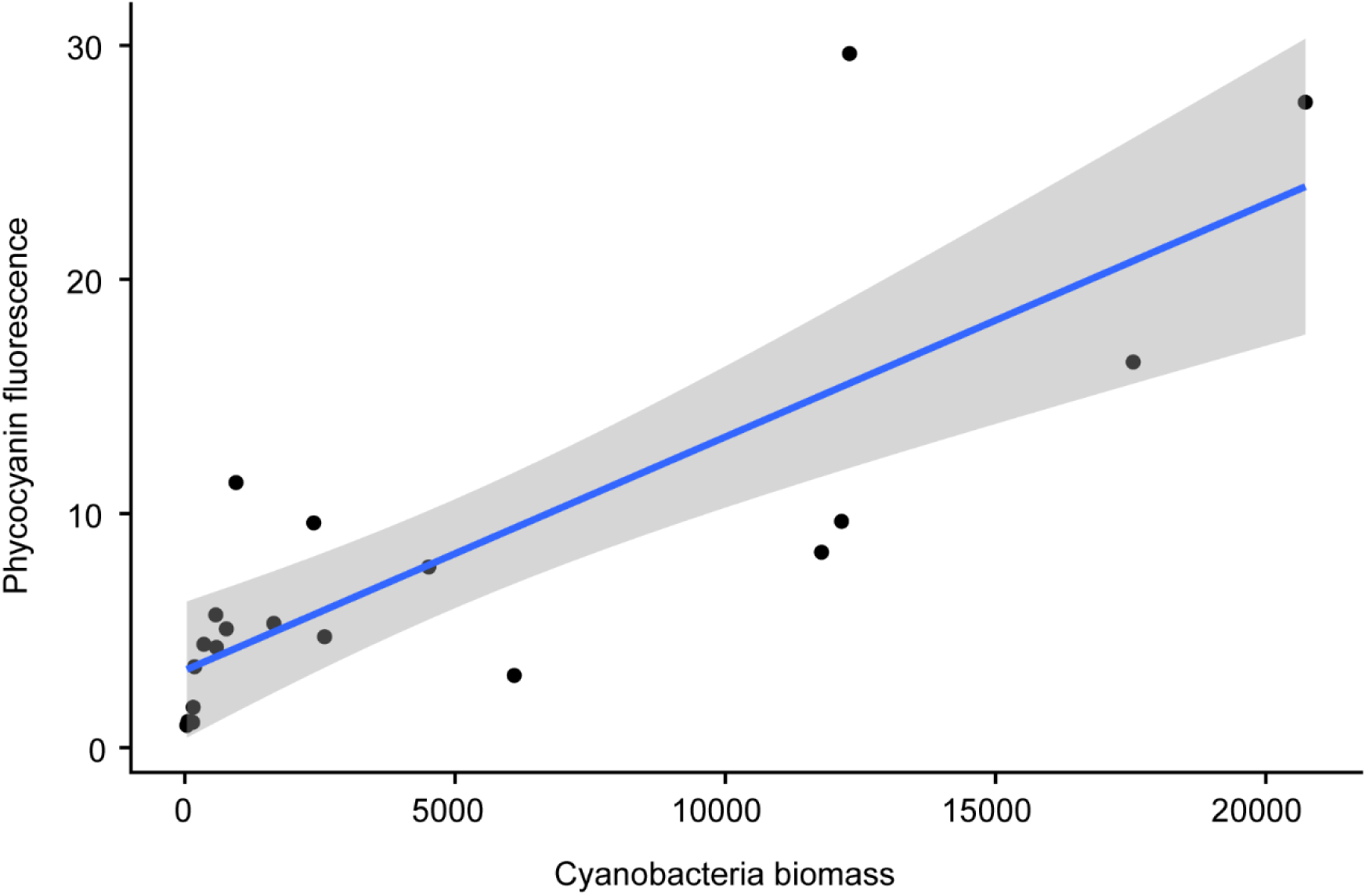
Buoy phycocyanin fluorescence plotted against cyanobacteria biomass for 2014- 2017. Water samples were collected at approximately 0.1m depth adjacent the buoy (slightly higher in the water column than the phycocyanin sensor – see main text) and also from the water treatment plant offtake which is located at approximately 3.0 m depth near the buoy. Blue line is the fitted regression while the grey area is the 95% confidence interval. Cyanobacteria biomass is in units of mg.m^-3^ and phycocyanin fluorescence has the unit relative fluorescence unit (RFU).

**Figure A2.**
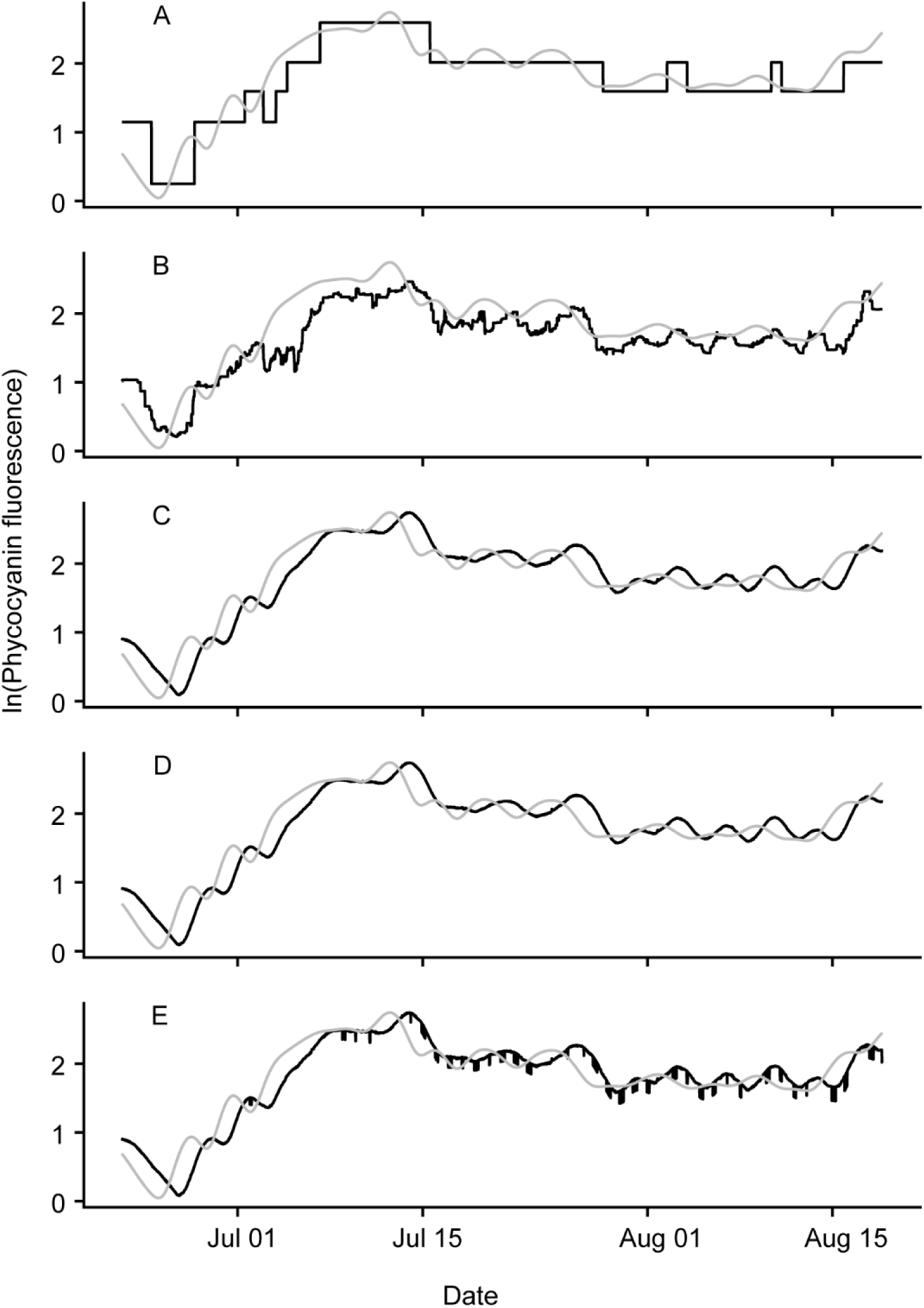
1 day forecast for 2014. A) decision tree (dt) B) Boosted decision tree (bdt) C) linear regression (lm) D) boosted linear regression (blm) E) non-linear model (nlm). Grey line is observed while black line is forecasted. Note that the unit for phycocyanin fluorescence is relative fluorescence unit (RFU).

**Figure A3.**
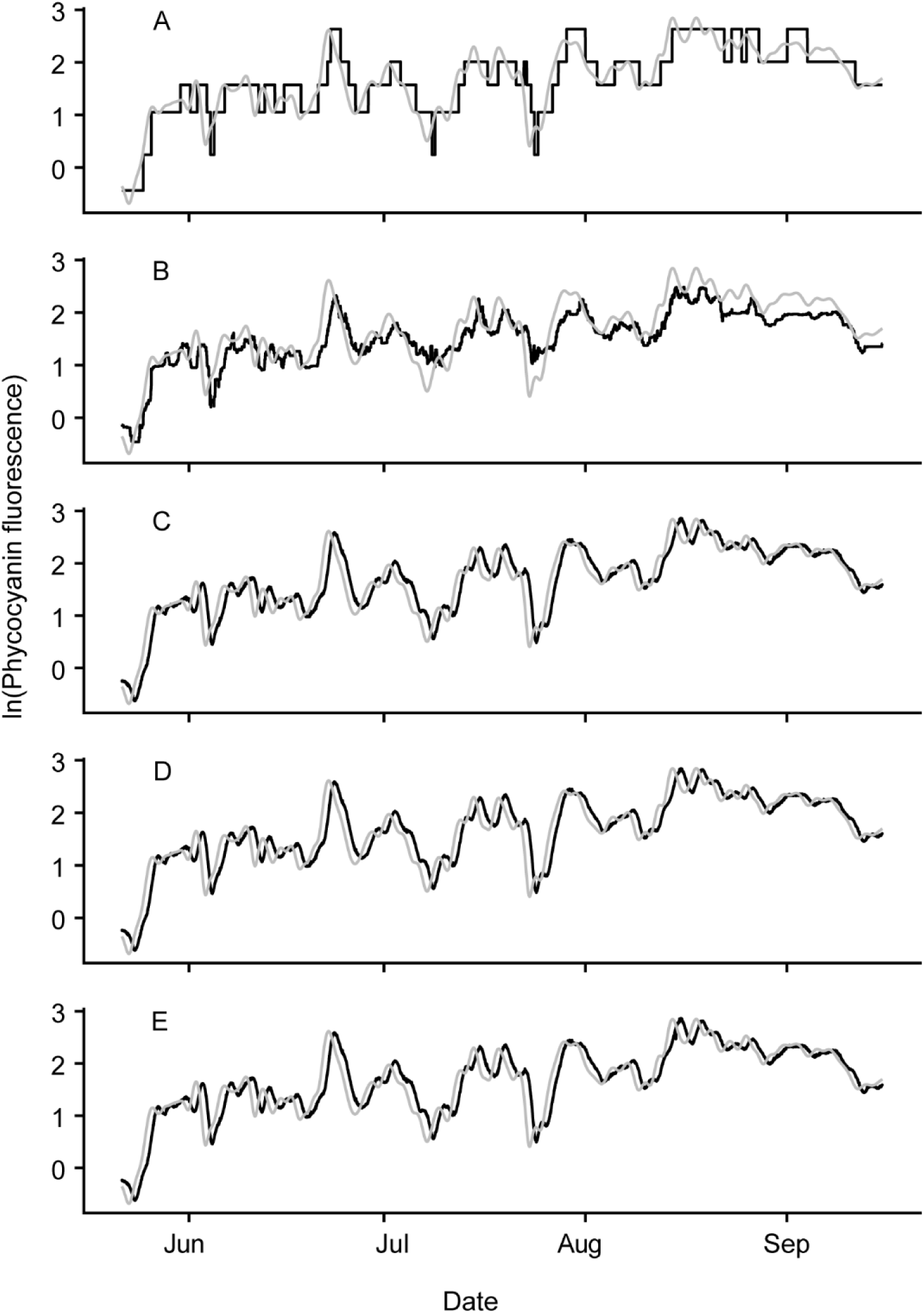
1 day forecast for 2015. A) decision tree (dt) B) Boosted decision tree (bdt) C) linear regression (lm) D) boosted linear regression (blm) E) non-linear model (nlm). Grey line is observed while black line is forecasted. Note that the unit for phycocyanin fluorescence is relative fluorescence unit (RFU).

**Figure A4.**
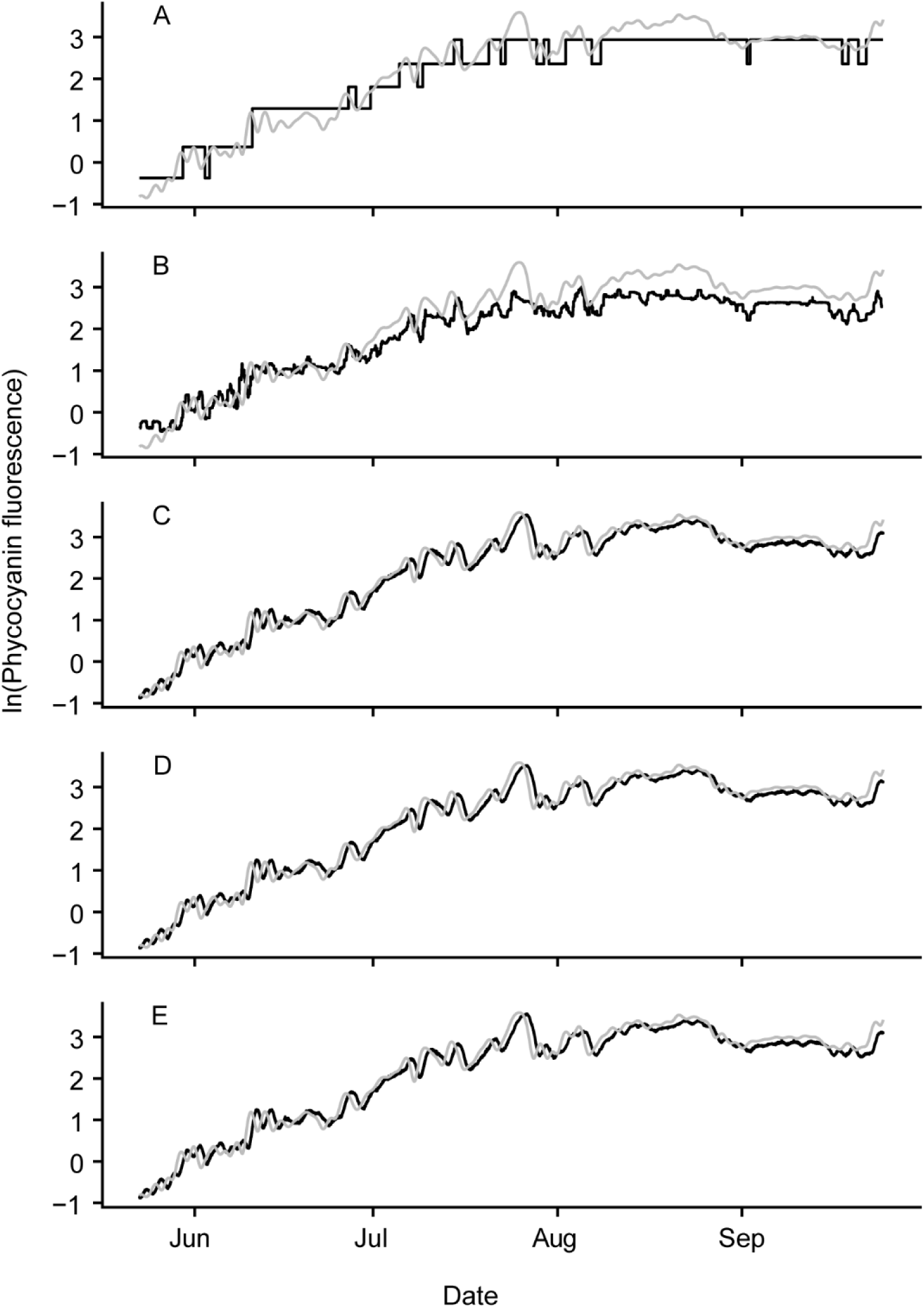
1 day forecast for 2016. A) decision tree (dt) B) Boosted decision tree (bdt) C) linear regression (lm) D) boosted linear regression (blm) E) non-linear model (nlm). Grey line is observed while black line is forecasted. Note that the unit for phycocyanin fluorescence is relative fluorescence unit (RFU).

**Figure A5.**
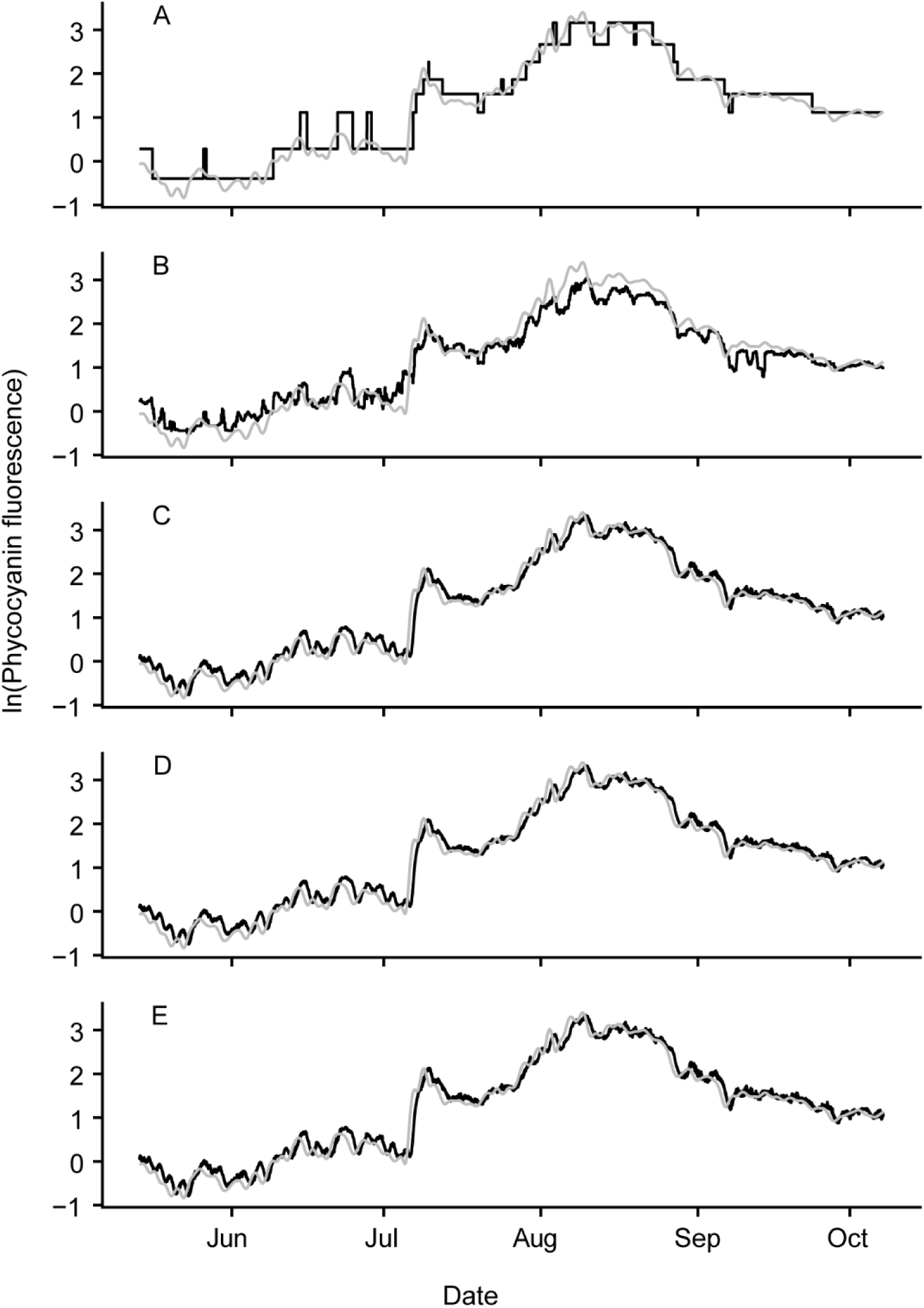
1 day forecast for 2017. A) decision tree (dt) B) Boosted decision tree (bdt) C) linear regression (lm) D) boosted linear regression (blm) E) non-linear model (nlm). Grey line is observed while black line is forecasted. Note that the unit for phycocyanin fluorescence is relative fluorescence unit (RFU).

**Figure A6.**
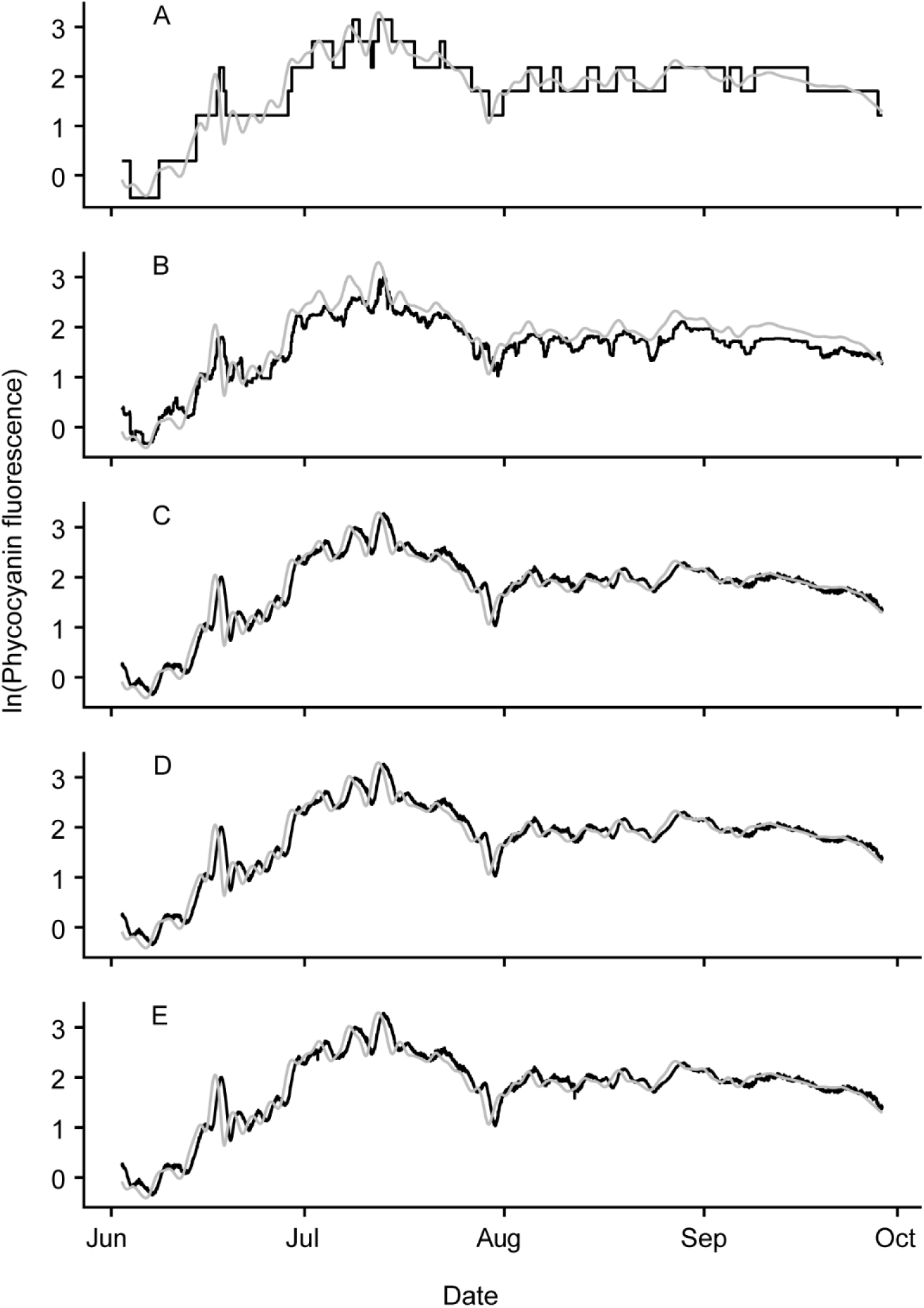
1 day forecast for 2018. A) decision tree (dt) B) Boosted decision tree (bdt) C) linear regression (lm) D) boosted linear regression (blm) E) non-linear model (nlm). Grey line is observed while black line is forecasted. Note that the unit for phycocyanin fluorescence is relative fluorescence unit (RFU).

**Figure A7.**
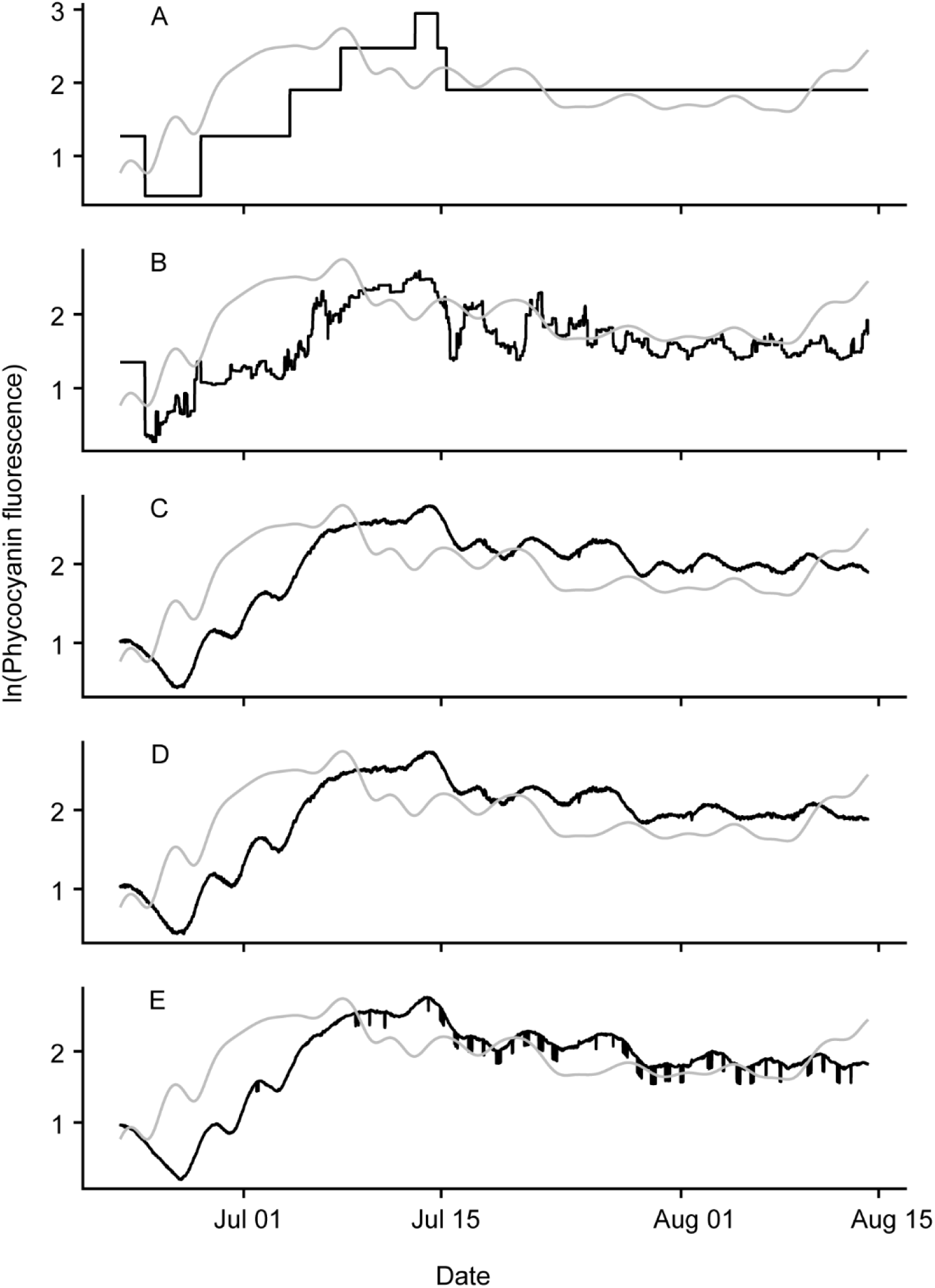
4 day forecast for 2014. A) decision tree (dt) B) Boosted decision tree (bdt) C) linear regression (lm) D) boosted linear regression (blm) E) non-linear model (nlm). Grey line is observed while black line is forecasted. Note that the unit for phycocyanin fluorescence is relative fluorescence unit (RFU).

**Figure A8.**
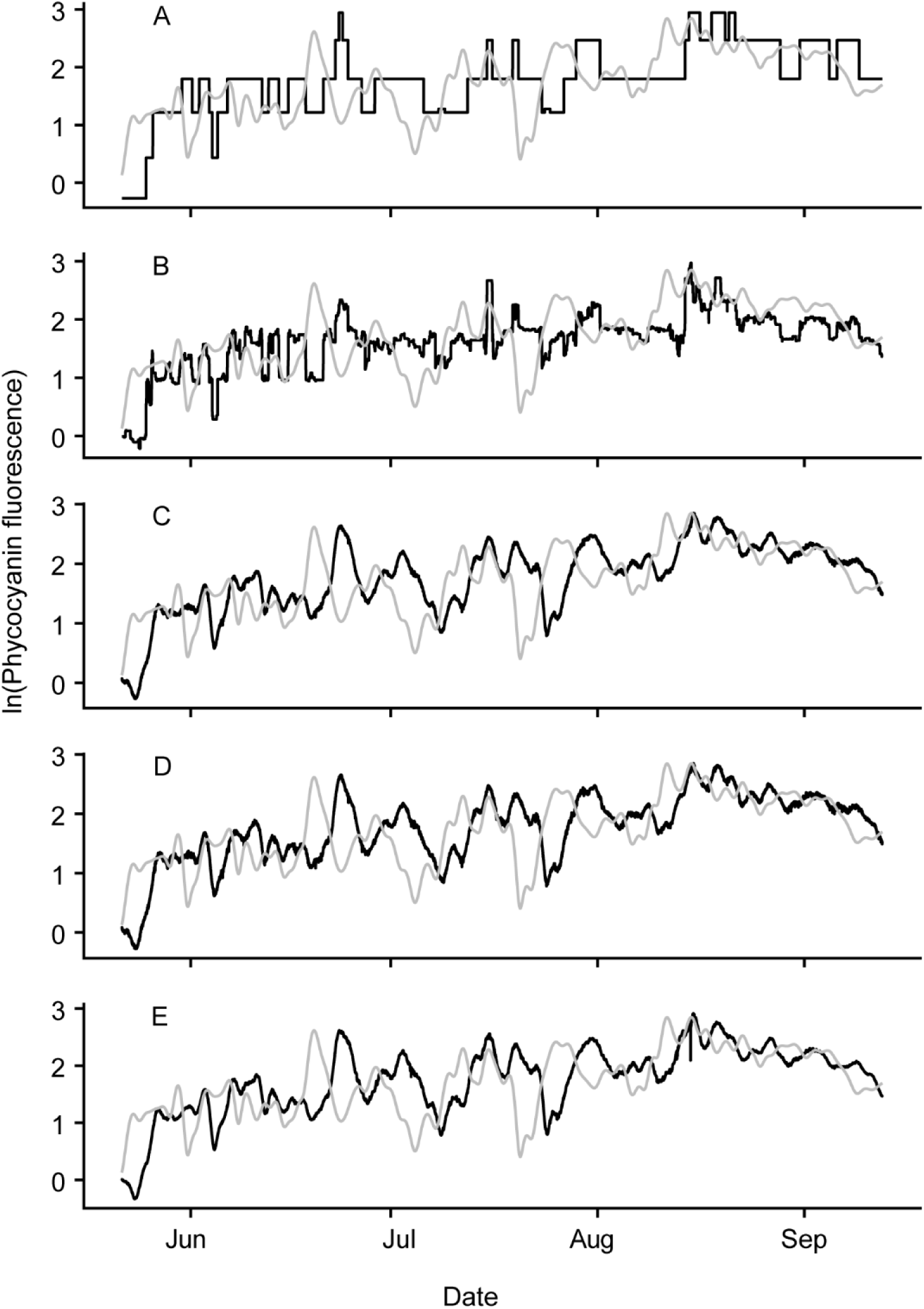
4 day forecast for 2015. A) decision tree (dt) B) Boosted decision tree (bdt) C) linear regression (lm) D) boosted linear regression (blm) E) non-linear model (nlm). Grey line is observed while black line is forecasted. Note that the unit for phycocyanin fluorescence is relative fluorescence unit (RFU).

**Figure A9.**
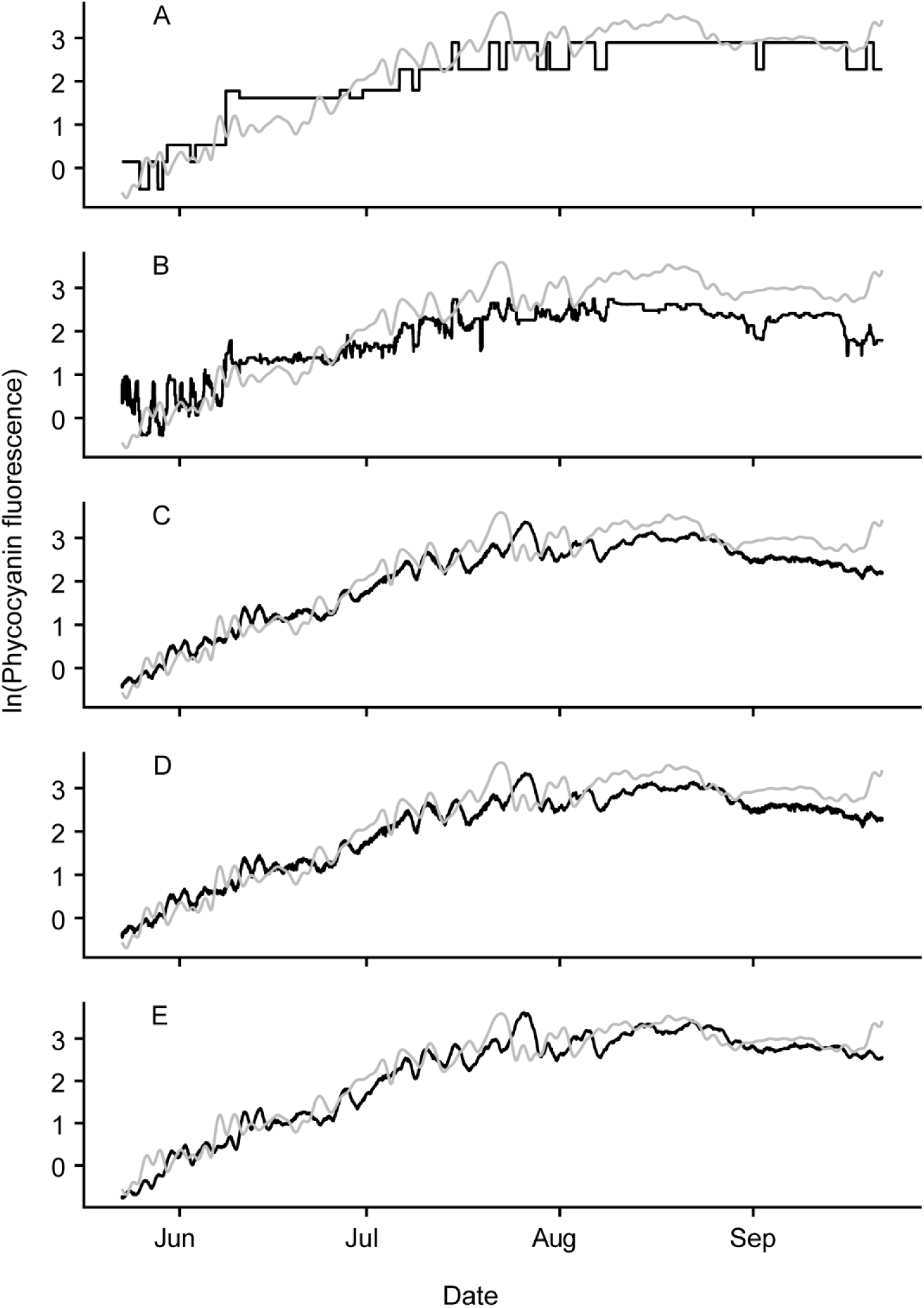
4 day forecast for 2016. A) decision tree (dt) B) Boosted decision tree (bdt) C) linear regression (lm) D) boosted linear regression (blm) E) non-linear model (nlm). Grey line is observed while black line is forecasted. Note that the unit for phycocyanin fluorescence is relative fluorescence unit (RFU).

**Figure A10.**
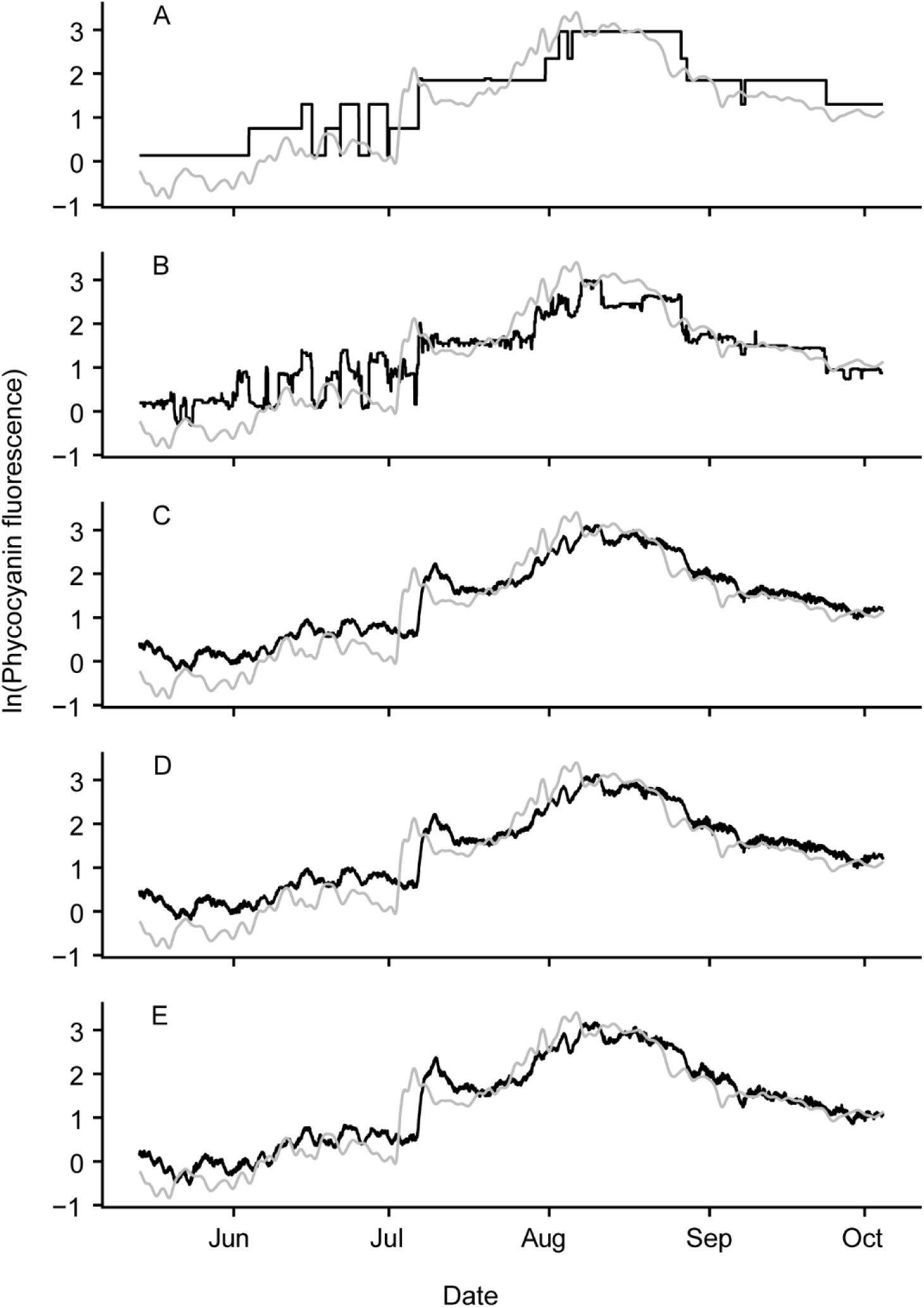
4 day forecast for 2017. A) decision tree (dt) B) Boosted decision tree (bdt) C) linear regression (lm) D) boosted linear regression (blm) E) non-linear model (nlm). Grey line is observed while black line is forecasted. Note that the unit for phycocyanin fluorescence is relative fluorescence unit (RFU).

**Figure A11.**
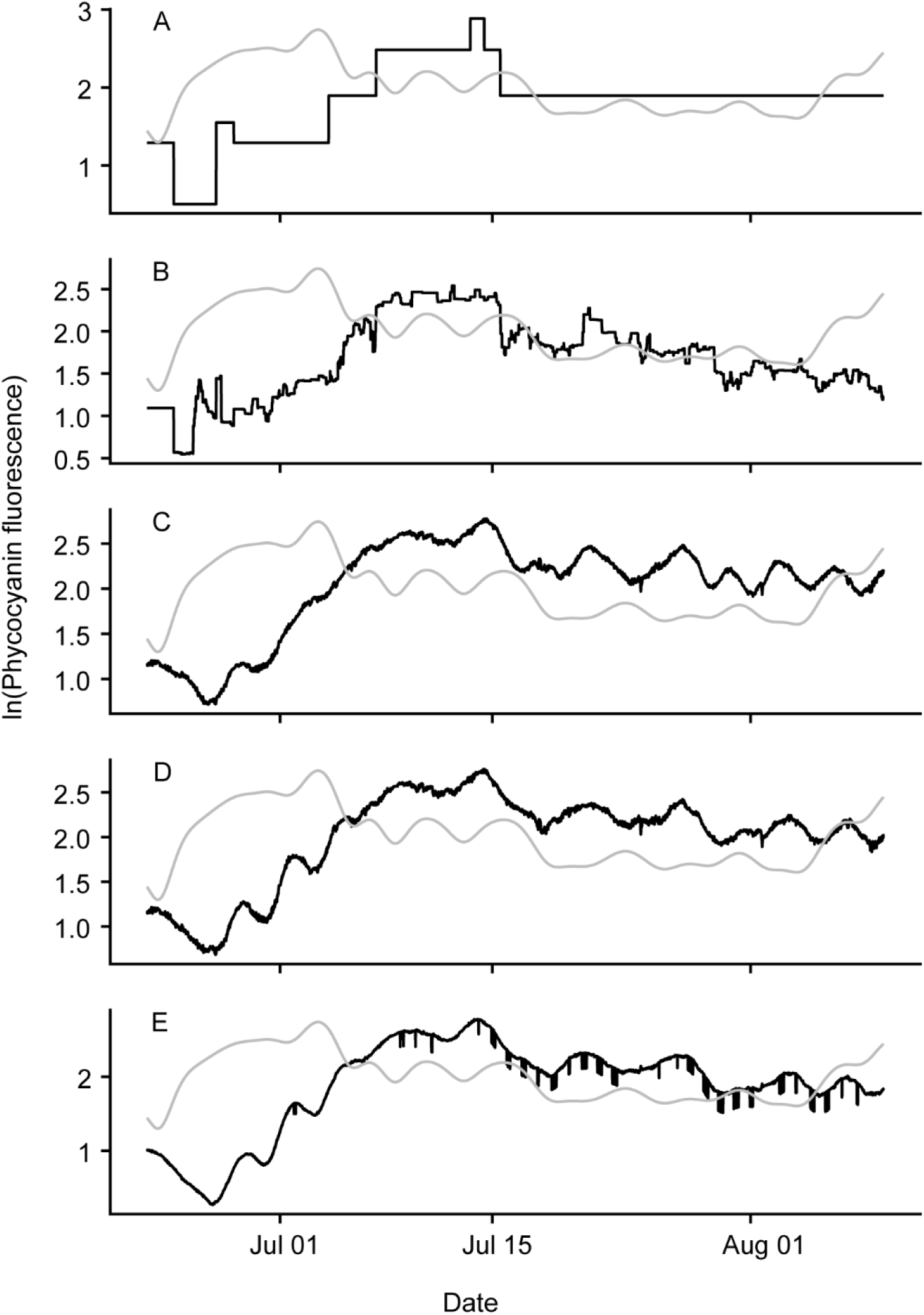
7 day forecast for 2014. A) decision tree (dt) B) Boosted decision tree (bdt) C) linear regression (lm) D) boosted linear regression (blm) E) non-linear model (nlm). Grey line is observed while black line is forecasted. Note that the unit for phycocyanin fluorescence is relative fluorescence unit (RFU).

**Figure A12.**
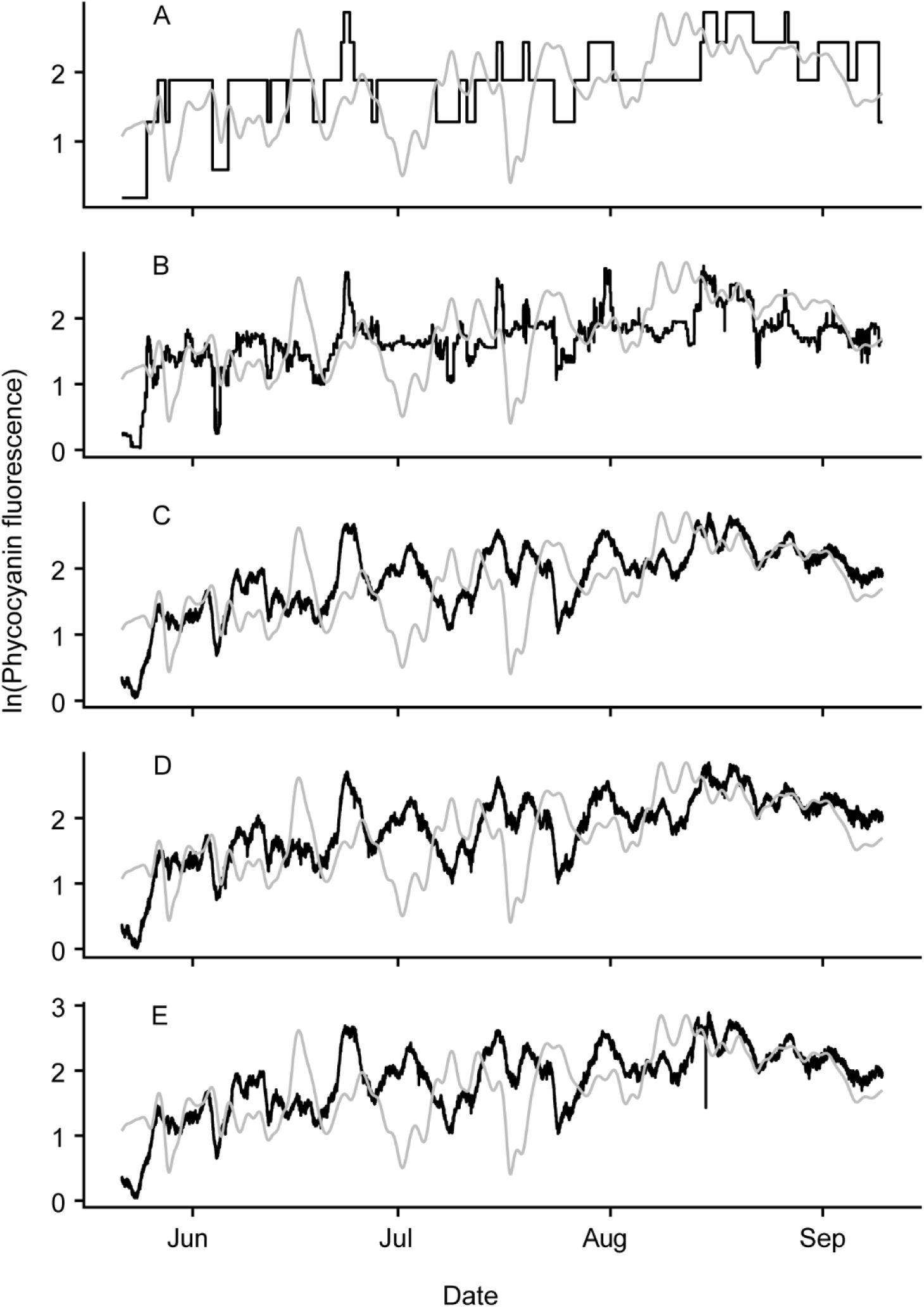
7 day forecast for 2015. A) decision tree (dt) B) Boosted decision tree (bdt) C) linear regression (lm) D) boosted linear regression (blm) E) non-linear model (nlm). Grey line is observed while black line is forecasted. Note that the unit for phycocyanin fluorescence is relative fluorescence unit (RFU).

**Figure A13.**
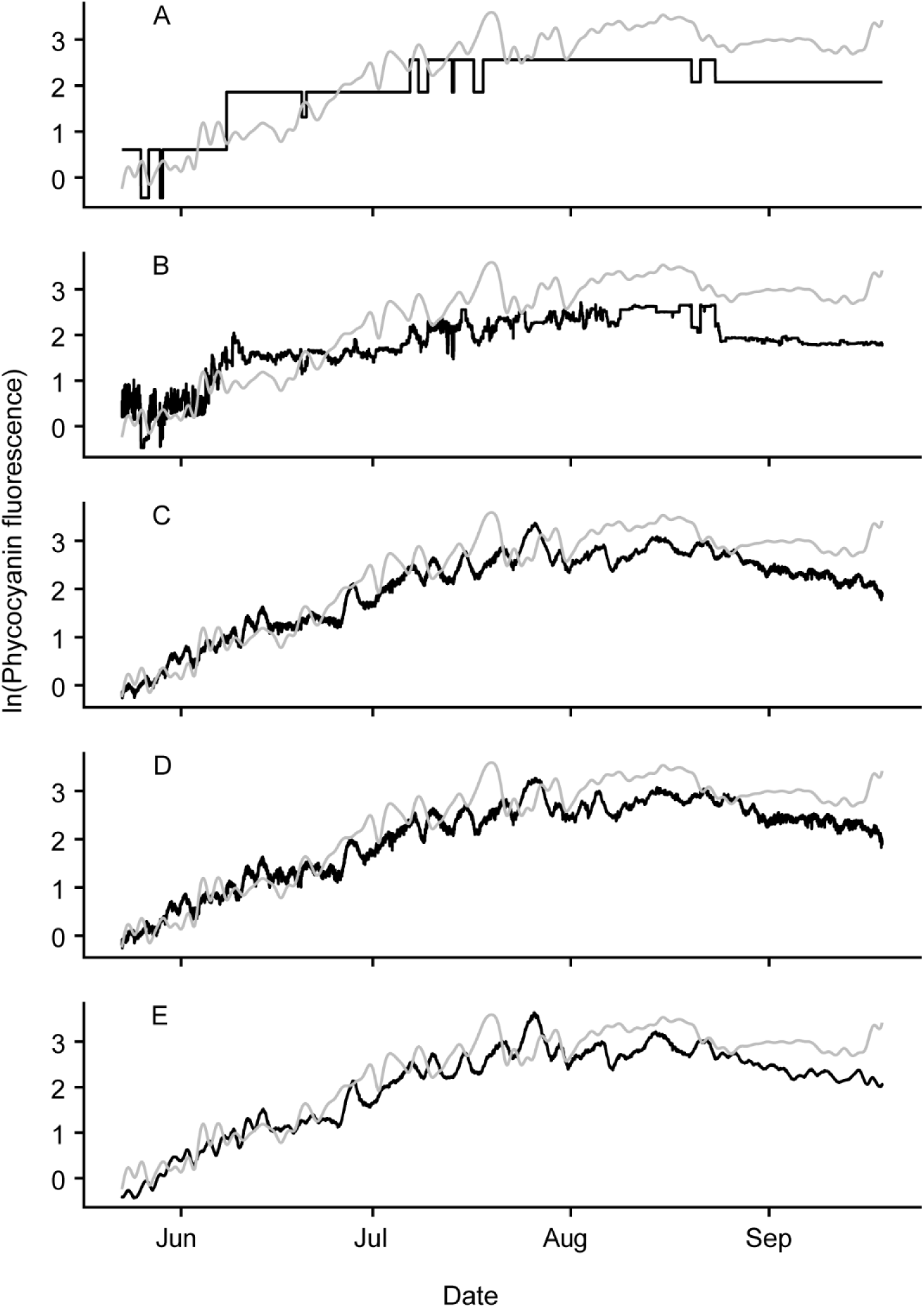
7 day forecast for 2016. A) decision tree (dt) B) Boosted decision tree (bdt) C) linear regression (lm) D) boosted linear regression (blm) E) non-linear model (nlm). Grey line is observed while black line is forecasted. Note that the unit for phycocyanin fluorescence is relative fluorescence unit (RFU).

**Figure A14.**
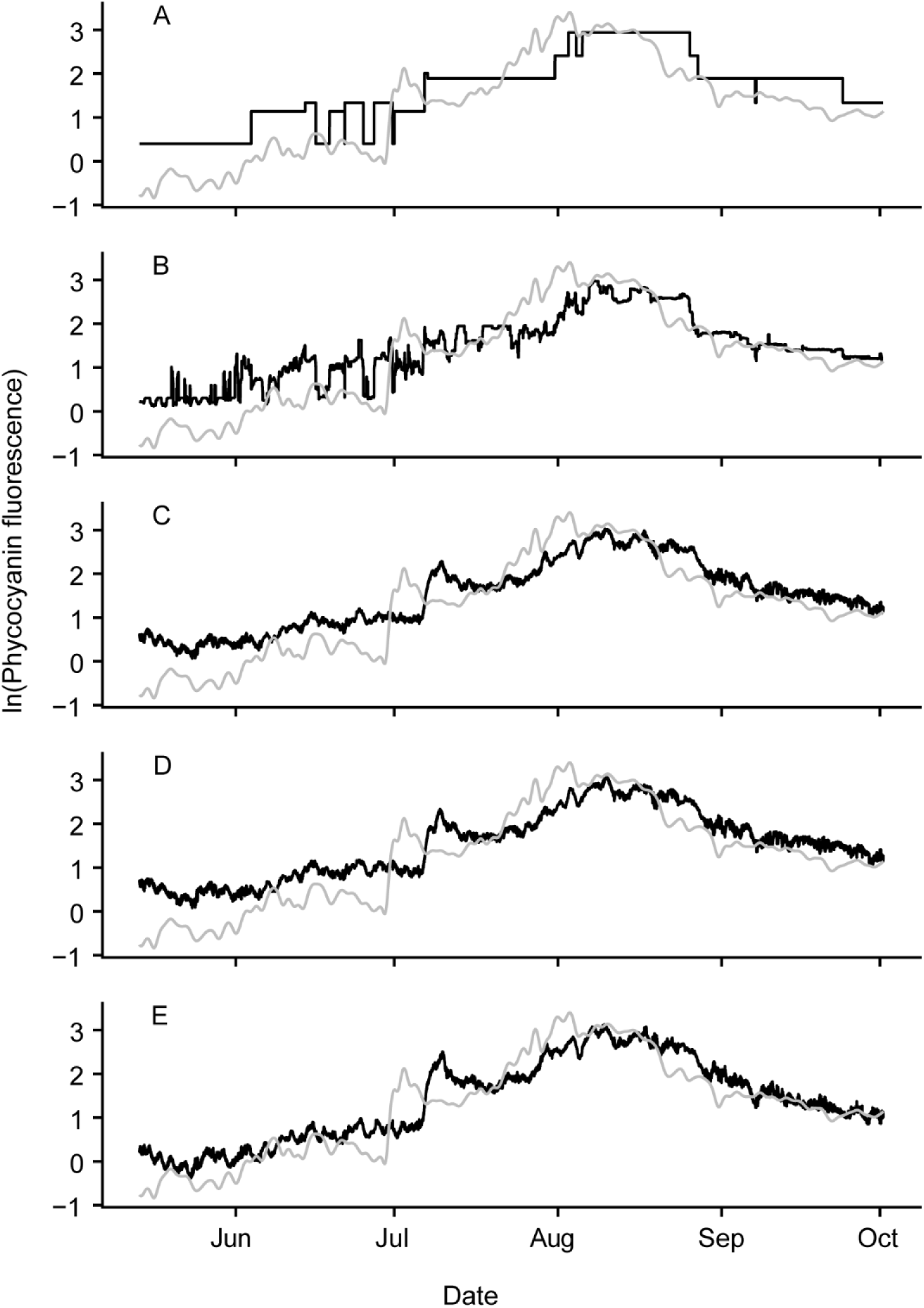
7 day forecast for 2017. A) decision tree (dt) B) Boosted decision tree (bdt) C) linear regression (lm) D) boosted linear regression (blm) E) non-linear model (nlm). Grey line is observed while black line is forecasted. Note that the unit for phycocyanin fluorescence is relative fluorescence unit (RFU).

**Figure A15.**
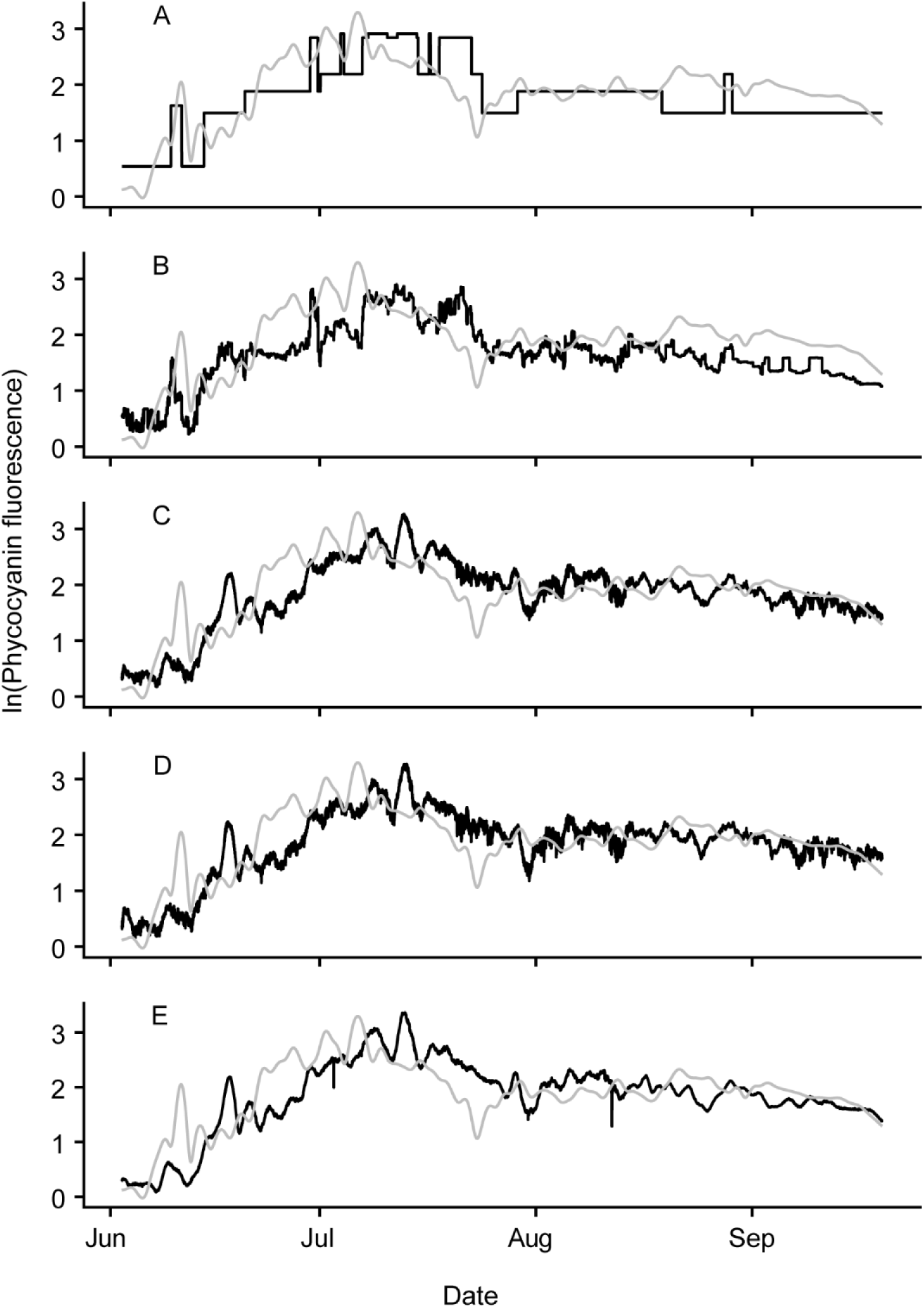
7 day forecast for 2018. A) decision tree (dt) B) Boosted decision tree (bdt) C) linear regression (lm) D) boosted linear regression (blm) E) non-linear model (nlm). Grey line is observed while black line is forecasted. Note that the unit for phycocyanin fluorescence is relative fluorescence unit (RFU).

**Figure A16.**
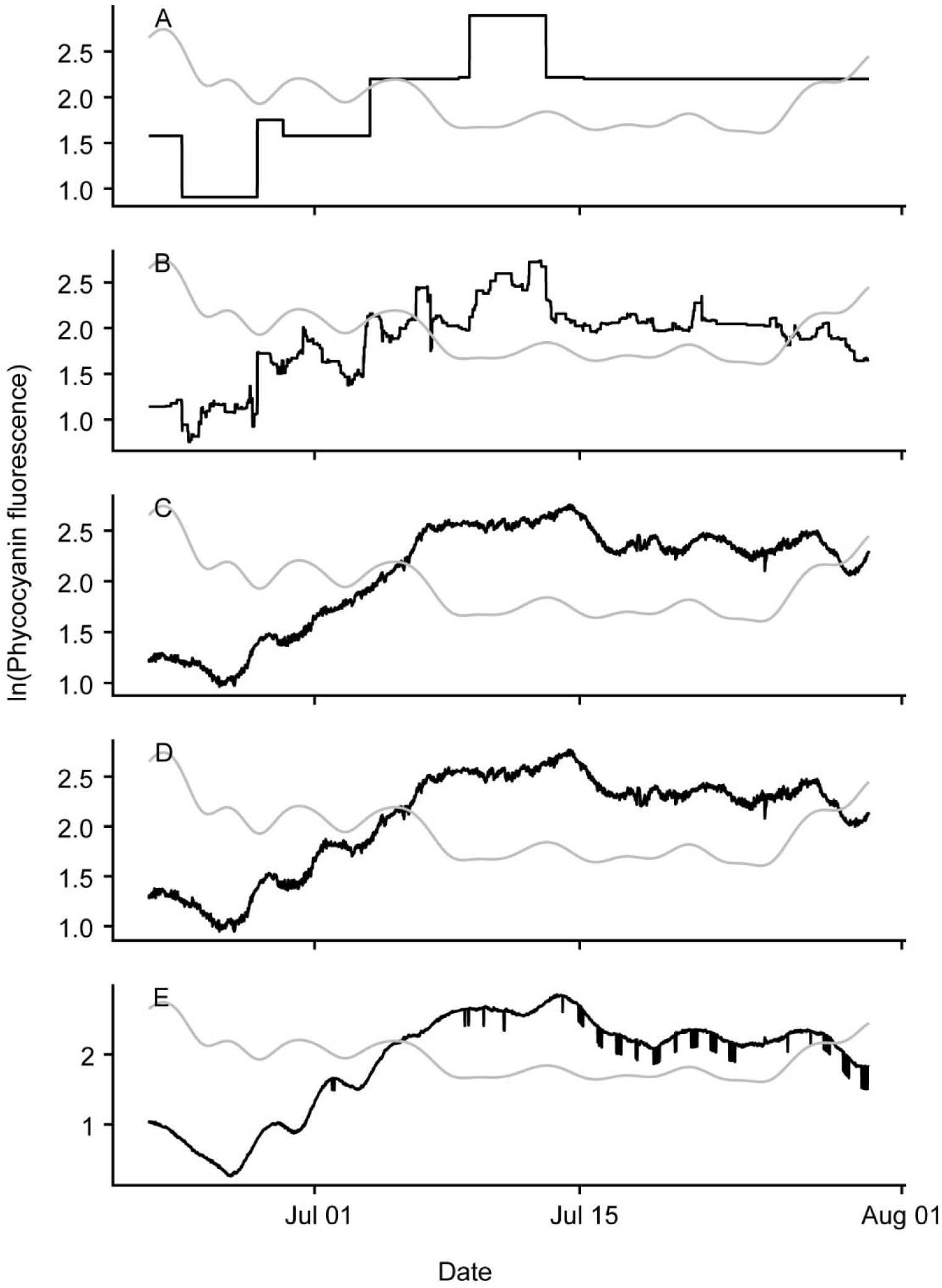
14 day forecast for 2014. A) decision tree (dt) B) Boosted decision tree (bdt) C) linear regression (lm) D) boosted linear regression (blm) E) non-linear model (nlm). Grey line is observed while black line is forecasted. Note that the unit for phycocyanin fluorescence is relative fluorescence unit (RFU).

**Figure A17.**
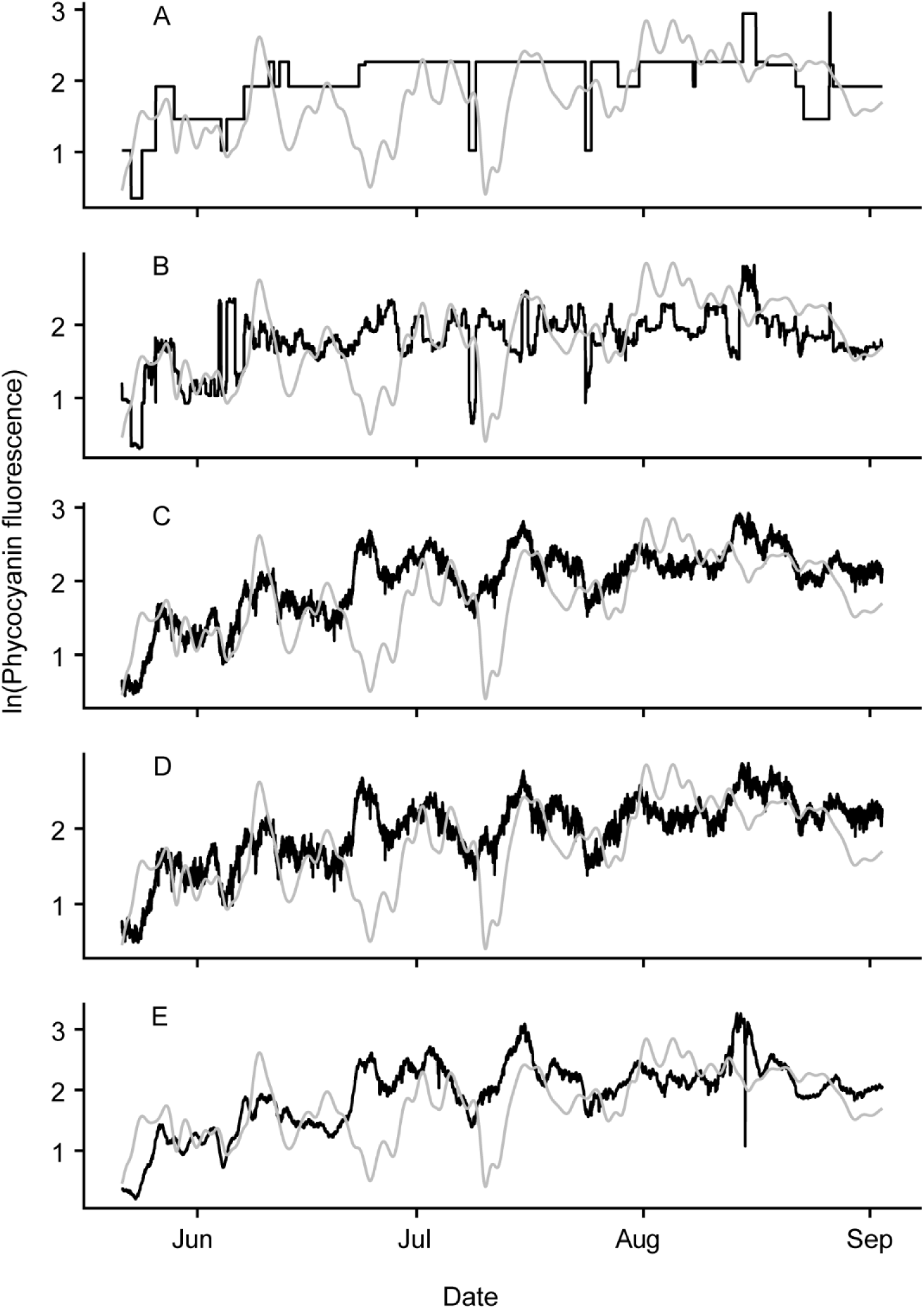
14 day forecast for 2015. A) decision tree (dt) B) Boosted decision tree (bdt) C) linear regression (lm) D) boosted linear regression (blm) E) non-linear model (nlm). Grey line is observed while black line is forecasted. Note that the unit for phycocyanin fluorescence is relative fluorescence unit (RFU).

**Figure A18.**
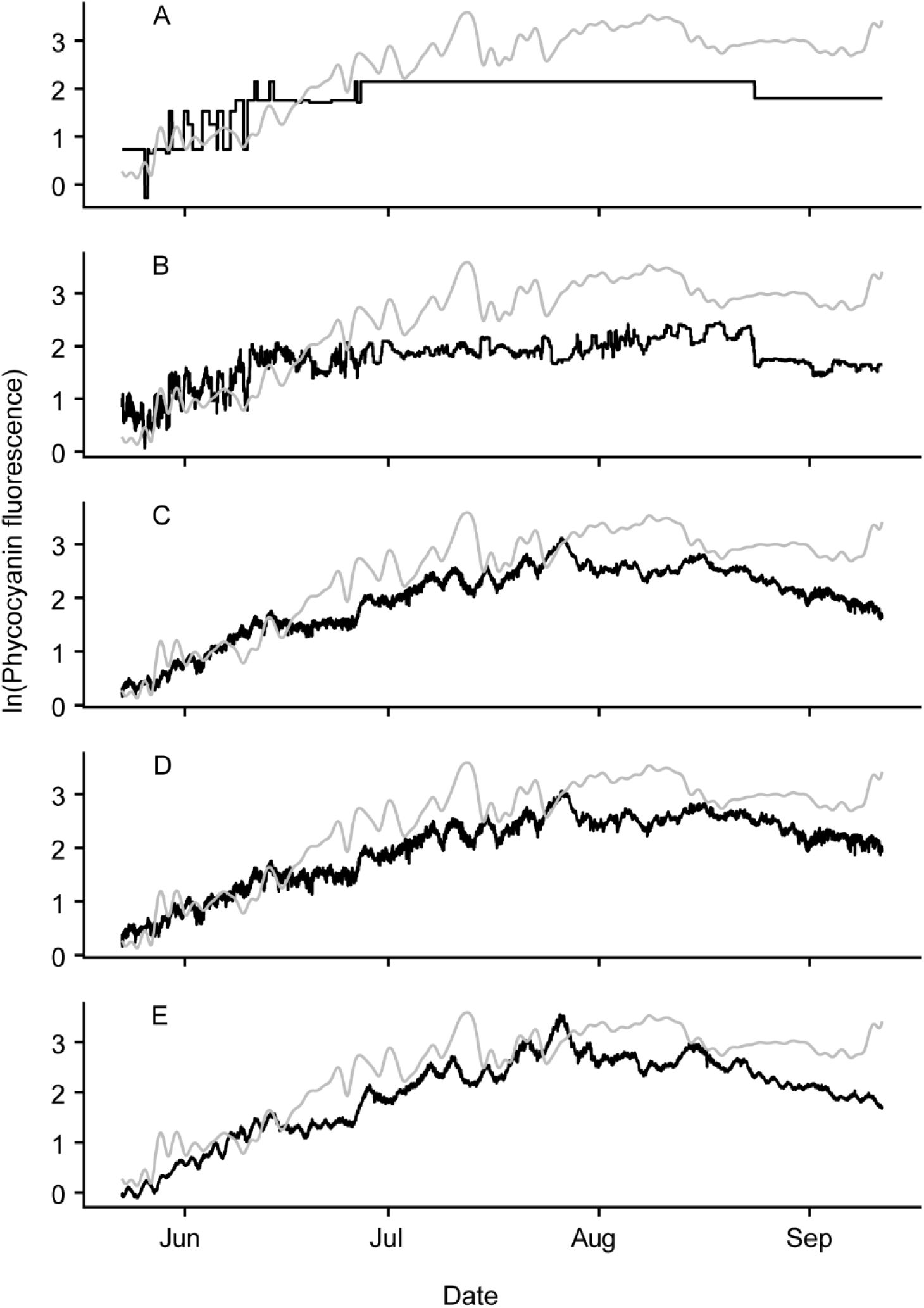
14 day forecast for 2016. A) decision tree (dt) B) Boosted decision tree (bdt) C) linear regression (lm) D) boosted linear regression (blm) E) non-linear model (nlm). Grey line is observed while black line is forecasted. Note that the unit for phycocyanin fluorescence is relative fluorescence unit (RFU).

**Figure A19.**
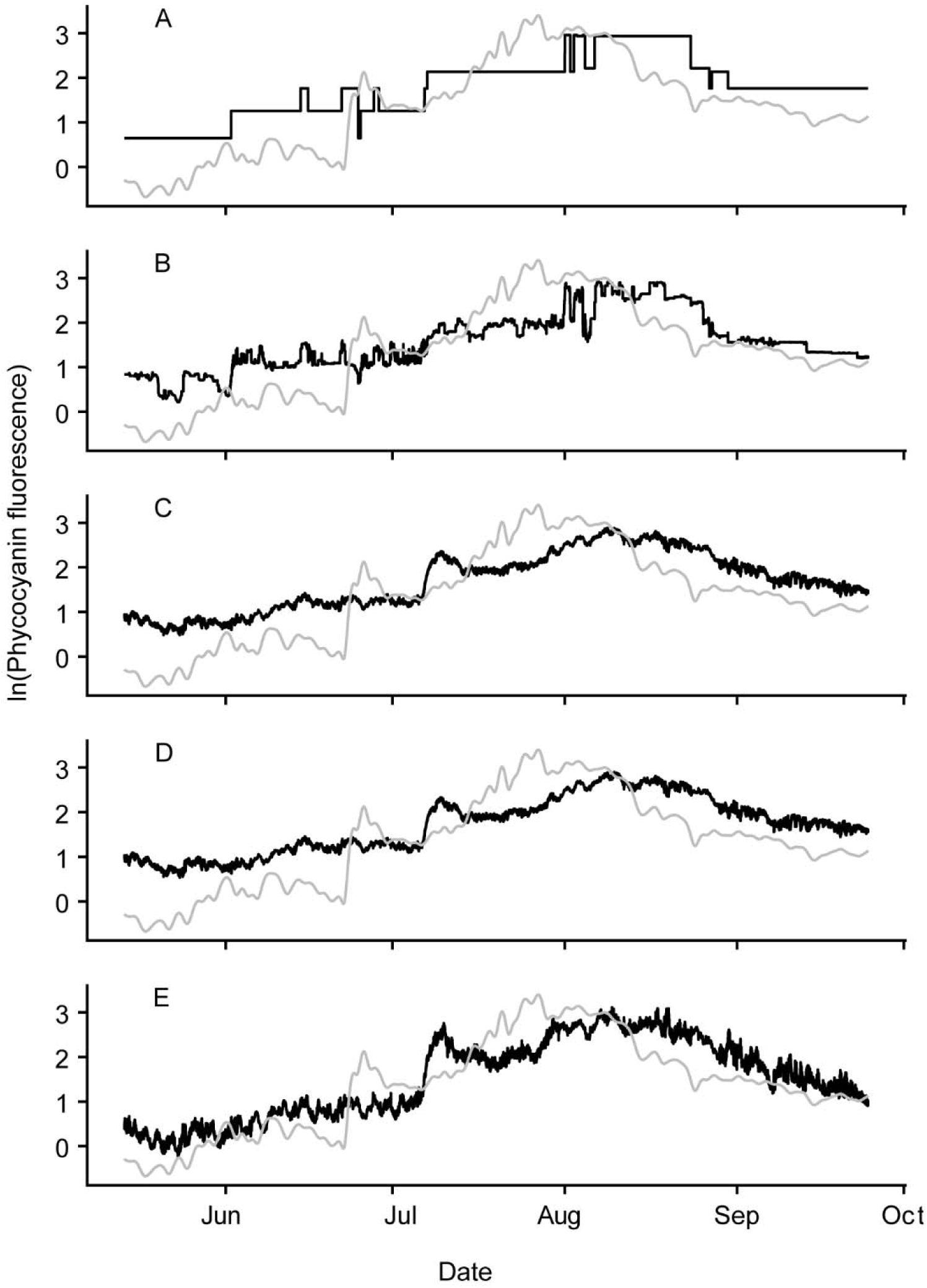
14 day forecast for 2017. A) decision tree (dt) B) Boosted decision tree (bdt) C) linear regression (lm) D) boosted linear regression (blm) E) non-linear model (nlm). Grey line is observed while black line is forecasted. Note that the unit for phycocyanin fluorescence is relative fluorescence unit (RFU).

**Figure A20.**
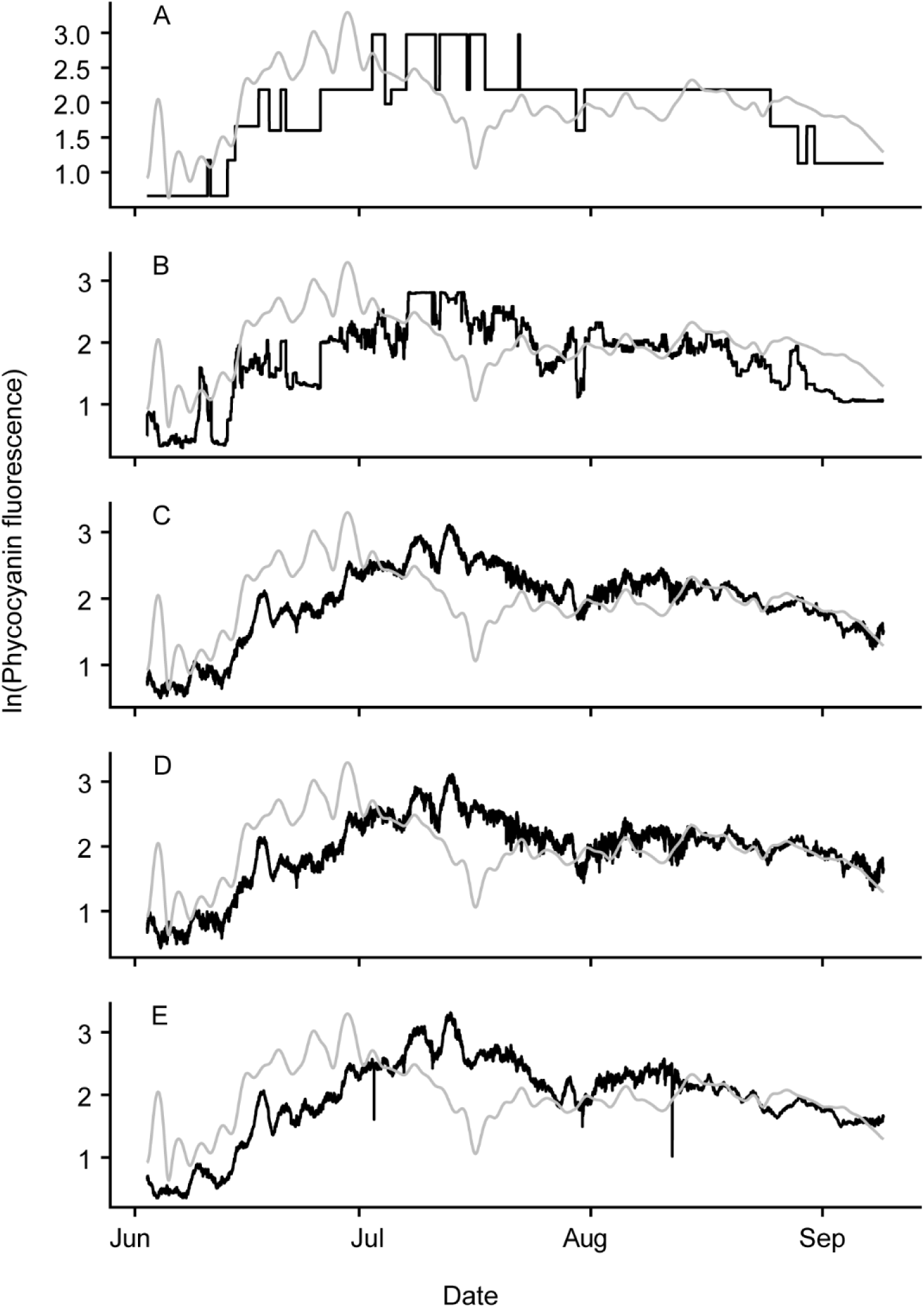
14 day forecast for 2018. A) decision tree (dt) B) Boosted decision tree (bdt) C) linear regression (lm) D) boosted linear regression (blm) E) non-linear model (nlm). Grey line is observed while black line is forecasted. Note that the unit for phycocyanin fluorescence is relative fluorescence unit (RFU).

## References

Chan, W.S., Recknagel, F., Cao, H., Park, H.-D., 2007. Elucidation and short-term forecasting of microcystin concentrations in Lake Suwa (Japan) by means of artificial neural networks and evolutionary algorithms. Water Research 41, 2247–2255.

Chen, T., He, T., Benesty, M., Khotilovich, V., Tang, Y., Cho, H., Chen, K., Mitchell, R., Cano, I., Zhou, T., Li, M., Xie, J., Lin, M., Geng, Y., Li, Y., 2018. xgboost: Extreme Gradient Boosting.

Dodds, W.K., Bouska, W.W., Eitzmann, J.L., Pilger, T.J., Pitts, K.L., Riley, A.J., Schloesser, J.T., Thornbrugh, D.J., 2009. Eutrophication of US freshwaters: analysis of potential economic damages. Enviromental Science and Technology 43(1), 12–19.

Fernández, J.A., Muñiz, C.D., Nieto, P.G., de Cos Juez, F., Lasheras, F.S., Roqueñí, M., 2013. Forecasting the cyanotoxins presence in fresh waters: A new model based on genetic algorithms combined with the MARS technique. Ecological Engineering 53, 68–78.

Hall, R.I., Leavitt, P.R., Quinlan, R., Dixit, A.S., Smol, J.P., 1999. Effects of agriculture, urbanization, and climate on water quality in the northern Great Plains. Limnology and Oceanography 44, 739– 756.

Hamilton, D.P., Carey, C.C., Arvola, L., Arzberger, P., Brewer, C., Cole, J.J., Gaiser, E., Hanson, P.C., Ibelings, B.W., Jennings, E., 2015. A Global Lake Ecological Observatory Network (GLEON) for synthesising high-frequency sensor data for validation of deterministic ecological models. Inland Waters 5, 49–56.

Hammer, U.T., 1978. The saline lakes of Saskatchewan III. Chemical characterization. Internationale Revue der gesamten Hydrobiologie und Hydrographie 63, 311–335.

Hoagland, P., Anderson, D.M., Kaoru, Y., White, A.W., 2002. The economic effects of harmful algal blooms in the United States: estimates, assessment issues, and information needs. Estuaries 25, 819–837.

Hodges, C.M., Wood, S.A., Puddick, J., McBride, C.G., Hamilton, D.P., 2018. Sensor manufacturer, temperature, and cyanobacteria morphology affect phycocyanin fluorescence measurements. Environmental Science and Pollution Research. 25(2), 1079–1088.

Hudnell, H.K., 2010. The state of US freshwater harmful algal blooms assessments, policy and legislation. Toxicon 55, 1024–1034.

Kehoe, M.J., Chun, K.P., Baulch, H.M., 2015. Who Smells? Forecasting Taste and Odor in a Drinking Water Reservoir. Environmental Science and Technology 49, 10984–10992.

Kong, F., Ma, R., Gao, J., Wu, X., 2009. The theory and practice of prevention, forecast and warning on cyanobacteria bloom in Lake Taihu. Journal of Lake Sciences 3, 314–328.

Kotak, B.G., Zurawell, R.W., 2007. Cyanobacterial toxins in Canadian freshwaters: A review. Lake and Reservoir Management 23, 109–122.

Li, W., Qin, B., Zhu, G., 2014. Forecasting short-term cyanobacterial blooms in Lake Taihu, China, using a coupled hydrodynamic–algal biomass model. Ecohydrology 7, 794–802.

Mantzouki, E., Lürling, M., Fastner, J., de Senerpont Domis, L., Wilk-Woźniak, E., Koreivienė, J., Seelen, L., Teurlincx, S., Verstijnen, Y., Krztoń, W., 2018. Temperature effects explain continental scale distribution of cyanobacterial toxins. Toxins 10, 156.

Obenour, D.R., Gronewold, A.D., Stow, C.A., Scavia, D., 2014. Using a Bayesian hierarchical model to improve Lake Erie cyanobacteria bloom forecasts. Water Resources Research 50, 7847–7860.

Onderka, M., 2007. Correlations between several environmental factors affecting the bloom events of cyanobacteria in Liptovska Mara reservoir (Slovakia)—A simple regression model. Ecological Modelling 209, 412–416.

Persaud, A.D., Paterson, A.M., Dillon, P.J., Winter, J.G., Palmer, M., Somers, K.M., 2015. Forecasting cyanobacteria dominance in Canadian temperate lakes. Journal of Environmental Management 151, 343–352.

Pick, F.R., 2016. Blooming algae: a Canadian perspective on the rise of toxic cyanobacteria. Canadian Journal of Fisheries and Aquatic Sciences 73, 1149–1158.

R Core Team, 2017. R: A Language and Environment for Statistical Computing. R Foundation for Statistical Computing, Vienna, Austria.

Seltenrich, N., 2014. Keeping tabs on HABs: new tools for detecting, monitoring, and preventing harmful algal blooms. Environmental Health Perspectives 122(8), A206–A213.

Therneau, T., Atkinson, B., Ripley, B., 2017. rpart: Recursive Partitioning and Regression Trees.

Thomas, M.K., Fontana, S., Reyes, M., Kehoe, M., Pomati, F., 2018. The predictability of a lake phytoplankton community, over time-scales of hours to years. Ecology letters 21, 619–628.

Welk, A., Recknagel, F., Cao, H., Chan, W.-S., Talib, A., 2008. Rule-based agents for forecasting algal population dynamics in freshwater lakes discovered by hybrid evolutionary algorithms. Ecological Informatics 3, 46–54.

Wickham, H., 2016. ggplot2: Elegant Graphics for Data Analysis. Springer-Verlag New York.

Winslow, L.A., Read, J.S., Woolway, R., Brentrup, J., Leach, T., Zwart, J.A., Albers, S., Collinge, D., 2018. rLakeAnalyzer: Lake Physics Tools.

Wynne, T.T., Stumpf, R.P., Tomlinson, M.C., Fahnenstiel, G.L., Dyble, J., Schwab, D.J., Joshi, S.J., 2013. Evolution of a cyanobacterial bloom forecast system in western Lake Erie: Development and initial evaluation. Journal of Great Lakes Research 39, 90–99.

## References

Findlay, D. L., and H. J. Kling. 2001. Protocols for monitoring biodiversity: Phytoplankton in freshwaters. Ecological Monitoring and Assessment Network (EMAN), Environment Canada. Ecological Monitoring and Assessment Network (Environment Canada).

Rott E. 1981. Some results from phytoplankton counting intercalibrations. Schweiz. Z. Hydrobiologia, 43, 43–62.

Vollenweider R.A. 1968. Scientific fundamentals of the eutrophication of lakes and flowing waters, with particular reference to nitrogen and phosphorus as factors in eutrophication. Technical Report, Organization for Economic Cooperation and Development, Paris, 27, 1–182.

